# AMPK Signalling Rewires WWP1 mediated RUNX1 Ubiquitination in CML Progression

**DOI:** 10.64898/2026.10.13.758885

**Authors:** Abhayananda Behera, Gorantla Sri Charitha, Maddimadugu Elina Rani, Nagaraj Velugonda, Dorababu Patchva, Sadasivudu Gundeti, Rama Krishna Kancha, Aramati Bindu Madhava Reddy

**Affiliations:** Department of Animal Biology, School of Life Sciences, University of Hyderabad, Hyderabad, 500046-India; Department of Medical Oncology, Nizam’s Institute of Medical Sciences, Hyderabad, 500082-India; Department of Pharmacology, Apollo Institute of Medical Sciences and Research, Jubilee Hills, Hyderabad, 500033-India; Molecular Medicine and Therapeutics Laboratory, CPMB, Osmania University, Hyderabad-500007, India

**Author notes:** To whom correspondence should be addressed Aramati Bindu Madhava Reddy, Department of Animal Biology, School of Life Sciences, University of Hyderabad, Hyderabad, 500046-India.

**Keywords:** WWP1, RUNX1, AMPK, CML, metformin, ubiquitination, phosphorylation

## Abstract

Despite the remarkable success of BCR:ABL1-targeted tyrosine kinase inhibitors, disease progression and therapeutic resistance remain major challenges in chronic myeloid leukemia (CML), highlighting the need to identify additional molecular mechanisms that sustain leukemic survival. In this study, we identify the HECT-type E3 ubiquitin ligase WWP1 as a previously unrecognized regulator of RUNX1 and define a signaling network linking ubiquitination to AMP-activated protein kinase (AMPK)-dependent phosphorylation in CML. We demonstrate that WWP1 directly associates with the PPxY motif of RUNX1 and catalyzes its K63-linked polyubiquitination, thereby enhancing protein stability and promoting leukemic cell growth. Disruption of WWP1, either by genetic silencing or pharmacological inhibition, diminished RUNX1 ubiquitination, impaired proliferation, induced G2/M cell-cycle arrest, and triggered apoptosis. Mechanistic investigations further revealed that WWP1 is a direct substrate of AMPK. Phosphorylation of WWP1 at Thr692 by AMPK promoted its lysosomal degradation, thereby limiting WWP1-dependent ubiquitination of RUNX1. In parallel, AMPK-mediated phosphorylation of RUNX1 reduced its leukemogenic activity, collectively shifting the signaling balance toward growth suppression. Activation of AMPK with metformin reproduced these molecular effects in leukemia cell lines, primary CML patient-derived cells, and xenograft tumors, supporting the physiological relevance of this pathway. Together, our findings establish an AMPK-WWP1-RUNX1 signaling axis that integrates phosphorylation and non-proteolytic ubiquitination to regulate RUNX1 function and CML progression. These results identify WWP1 as a potential therapeutic vulnerability and provide a mechanistic rationale for targeting the AMPK-WWP1 pathway in CML.

## Introduction

The runt-related transcription factor RUNX1, a member of the RUNX family, regulates hematopoiesis by orchestrating lineage commitment and differentiation. Its dysregulation is strongly linked to hematological malignancies, including CML [1, 2]. In leukemia, chromosomal rearrangements involving RUNX1 fusion with numerous partners, such as RUNX1-ETO [3], RUNX1-EVI1 [4], RUNX1-ETV6 [5], and RUNX1-T1 [6], are often reported, resulting in cell fate decisions and inhibiting lineage commitment [7, 8]. The stability and activity of RUNX1 are governed by posttranslational modifications (PTMs), notably ubiquitination and phosphorylation [9]. However, the molecular cues modulating RUNX1 turnover and function in leukemic contexts remain poorly understood.

Several known E3 ubiquitin ligases, such as APC [10], RNF38 [11], CHIP [12] and STUB1 [13] have been implicated in regulating RUNX1, primarily during development and erythroid lineage differentiation. However, their potential roles as oncogenes or tumor suppressors in leukemia remain largely unexplored. Notably, an AlphaScreen assay identified nine E3 ubiquitin ligases that interact with RUNX1, among which WWP1 (WW domain-containing E3 ubiquitin protein ligase 1), a HECT-type E3 ubiquitin ligase, emerged as a top candidate [13, 14]. WWP1 has been extensively studied in the context of solid tumors, where it promotes tumor growth and progression by targeting substrates involved in proliferation and survival pathways [15]. More recently, WWP1 has also been implicated in hematologic malignancies, including acute myeloid leukemia (AML) [16–18]. However, its role in CML remains unexplored.

Phosphorylation often primes or modulates ubiquitination, and the interplay between these PTMs is crucial for fine-tuning transcription factor stability [19]. RUNX1 phosphorylation has been demonstrated to increase transcriptional activity while also promoting degradation [9, 20]. For example, ERK-mediated phosphorylation of serine and threonine residues S249, S266, S276, and T273 activates RUNX1 [21], whereas CDK1/2/6 phosphorylation at identical locations marks it for APC-mediated degradation [10]. Our previous study demonstrated that AMP-activated protein kinase (AMPK), phosphorylates RUNX1 at serine 94, causing it to be sequestered in the cytoplasm via interaction with STAT3 [20, 22]. Interestingly, phosphorylated RUNX1 exerts an antileukemic effect in CML, suggesting a distinct axis of control.

In this study, we uncovered a dual regulatory mechanism involving WWP1-mediated K63-linked polyubiquitination of RUNX1 and WWP1 phosphorylation via AMPK. We demonstrated that WWP1 enhances leukemic proliferation via ubiquitination-mediated stabilization of RUNX1, whereas AMPK counters this effect by phosphorylating both WWP1 and RUNX1, resulting in WWP1 degradation and RUNX1 stability. These findings revealed an unidentified WWP1-AMPK-RUNX1 regulatory axis with important implications for the etiology of CML. Given the lack of clinically approved RUNX1-targeted therapies, our work provides a promising foundation for therapeutic strategies to modulate this regulatory pathway in leukemia.

## Materials and Methods

### In-silico analysis

A series of publicly available databases was employed for insilico analysis. RNAseq datasets from Cancer Cell Line Encyclopedia (CCLE) database (https://depmap.org/portal/) were used for WWP1 and RUNX1 expression analysis. Structural information for both WWP1 and RUNX1 was obtained from UniProt (https://www.uniprot.org/). Ubiquitination sites for RUNX1 were predicted using UbiBrowser 2.0 (https://ubibrowser.bio-it.cn/ubibrowser_v3/home/index). The mRNA expression levels of WWP1 and RUNX1 in AML cancer patients from TCGA were evaluated using the UALCAN database (http://ualcan.path.uab.edu/analysis.html), while their correlation was assessed via Pearson analysis using GEPIA (http://gepia.cancerpku.cn/detail.php?gene=&clicktag=expdiy). Overall Survival (OV) analysis was performed using the Kaplan-Meier (KM) plotter (https://kmplot.com/analysis/index.php?p=service&cancer=aml). To predict phosphorylation motifs, PhosphoMotif Finder (http://www.hprd.org/PhosphoMotif_finder/), and their significance scores were identified by the NetPhos 3.1 tool (http://www.cbs.dtu.dk/services/NetPhos/). All the protein sequences obtained from the NCBI human database (https://www.ncbi.nlm.nih.gov/) are used for sequence alignment by ClustalW2 (http://www.ebi.ac.uk/Tools/clustalw2/index.html).

### Isolation of PBMCs from CML patients

Blood samples from healthy individuals and patients were collected at Nizam’s Institute of Medical Sciences, Hyderabad, after obtaining approval from the Institutional Ethics Committee and written informed consent from all participants, in accordance with the approved guidelines (EC/NIMS/2350/2019). Approximately 6 mL of peripheral blood was obtained from each CML patient and from age-matched healthy individuals serving as controls, using heparin-coated vacutainers (BD, San Jose, CA, USA). The clinical and phenotypic characteristics of patients and healthy donors are summarized in Supplementary Table 1. Isolation of PBMCs (Peripheral blood mononuclear cells) was done using Histopaque (Sigma) density gradient centrifugation.

### Cell culture and chemicals

Human chronic myeloid leukemia (K562, KCL22), Human AML (HL-60), Human T-ALL (Jurkat J6), Human acute monocytic leukemia (THP1), and Human embryonic kidney (HEK293T) cell lines used in the study were purchased from ATCC (Manassas, VA, USA). All cells were maintained in RPMI1640 medium except HEK293T cells, which were maintained in DMEM at 37 °C in 5% CO2, with culture medium as suggested in the ATCC data sheet. All cells were mycoplasma-free and authenticated by short tandem repeat profiling. Metformin, Dorsomorphin (compound C), Indole-3-Carbinol (I3C), Puromycin, and Cycloheximide were purchased from Sigma (USA). shRNAs targeting WWP1 and RUNX1 (Supplementary Table 2) were purchased from the shRNA library at IISC, Bangalore, India.

### Construction of plasmids

RUNX1 cDNA (NM_001754.4) was purchased from DNASU (USA) (HS CD00630733). pCDNA3.1-HA-WWP1 was a kind gift from Prasad Tammineni, University of Hyderabad, India. WWP1 WT, WWP1 T692A, and WWP1 T692D were generated by site-directed mutagenesis and sub-cloned into pGEX-4T1 by using vector-gene-specific primers (Supplementary Table 3). pCDNA6.2/EmGFP-Bsd-RUNX1 (#176974), ΔRUNX AAYA and ΔRUNX AD were subcloned into pcDNA6.2/EmGFP-Bsd-RUNX1 vector, RUNX1 subcloned in pDsRed1-N1 by using the primers (Supplementary Table 2). pRK5-HA-WT Ub (#17608), K6 (#22900), K11 (#121152), K27 (#22902), K29 (#22903), K33 (#17607), K48 (#17605), K63 (#17606), and K0 (#17603), psPAX2 (# 12260) and pMD2.G (# 12259). were purchased from Addgene plasmid, USA [23]. WWP1 and catalytically inactive ΔWWP1-C890A in mammalian destination vector expressing Myc were generated using a gateway cloning system (Invitrogen), and pCDNA3.1-HA-K63R was a kind gift from Madika Suba Reddy, CDFD, Hyderabad, India.

### Transfection of plasmids

Cells were seeded in a 6-well plate and allowed to grow till 60-70% confluency. Then, the indicated plasmids were transfected with Lipofectamine-3000 reagents diluted in OPTI-MEM media as described earlier [20], and the cells were collected at the indicated time points for further analysis.

### shRNA transduction and generation of stable cell lines

HEK293T cells were transfected with either scrambled short hairpin RNA (shRNA), WWP1 shRNA, or RUNX1 shRNA as mentioned in the results description, along with viral packaging (psPAX2) vector and viral envelope vector (pMD2.G) by using Lipofectamine 3000 (Thermo Fisher, USA). At 24 h post-transfection, virus-containing media were collected, filtered through a 0.45 μm membrane filter, and the filtrate was used to transduce cells with 8 μg/ml polybrene (Sigma-Aldrich, USA) for 48 h. Post-transduction, the media were replaced with fresh complete media containing 1 μg/ml puromycin (Sigma, USA) until the selection was completed, compared with controls.

### PrestoBlue assay

Cell viability was measured according to the manufacturer’s instructions using the PrestoBlue assay. Briefly, 5×10^3^ K562 cells were seeded in 96-well plates with a total volume of 200 μl of RPMI medium treated with metformin and I3C at different concentrations on the same day. Following drug incubation at the specified time points, 10% v/v PrestoBlue (Thermo Fisher Scientific, A13261) was added to each well and incubated for 10 minutes at 37 °C; absorbance was then measured at 560-590 nm using a spectrophotometer.

### Western Blotting

Cells were harvested by adding 1X RIPA buffer containing protease and phosphatase inhibitor cocktail (Roche-04906837001, USA) and lysed by vortexing the cells for 30 minutes on ice, followed by centrifugation at 13000 rpm for 30 minutes. The supernatant was subjected to SDS-PAGE and transferred onto a nitrocellulose membrane according to standard protocols. The blots were probed with respective primary and secondary antibodies as listed in the supplementary table 4 [24].

### Nuclear and cytosolic extraction

The nuclear and cytosolic fractions were prepared as described previously [20]. In brief, the treated cells were harvested and centrifuged at 4000 rpm. Pellets were washed with 1X PBS, resuspended in buffer A, and incubated on ice for 15 minutes. After adding 10% NP-40 and vortexing for 10 minutes, samples were centrifuged at 8000 rpm for 10 minutes at 4 °C. The supernatant was collected as the cytosolic fraction. Buffer B was added to the pellet, which was then incubated on ice for 30 minutes with intermittent vortexing. After centrifugation at 14000 rpm for 20 minutes, the supernatant was collected as the nuclear fraction [20].

### Protein-protein docking

Protein-protein docking was performed to predict the potential interaction between WWP1 and RUNX1. The three-dimensional structure of human WWP1 was obtained from the Protein Data Bank (PDB ID: 6J1X), while the predicted structure of human RUNX1 was retrieved from the AlphaFold Protein Structure (AlphaFold ID: AF-Q01196-F1). Molecular docking was performed using HDOCK. The WW3 domain of WWP1 and the PPPY motif of RUNX1 were defined as regions of interest to investigate their potential involvement in the protein-protein interaction. The docking models generated by HDOCK were evaluated based on the docking scores and clustering parameters provided by the server, and the top-ranked model was selected for further analysis.

### Immunoprecipitation (IP)

For immunoprecipitation, 600-800 μg of protein lysate was incubated with the indicated antibodies in IP lysis buffer containing Tris-Cl, EDTA, NaCl, along with protease and phosphatase inhibitors for overnight binding at 4 ^0^C as described earlier [24]. The next day, the samples were incubated with 30 μg of equilibrated A/G agarose beads for 3 h at 4 °C. The bead complexes were washed 3 times with TBST buffer (Cell Signaling Technology) by centrifugation at 5000 rpm for 5 minutes. The bead-bound complexes were boiled in an IP lysis buffer containing 1X SDS loading buffer and analyzed for SDS-PAGE and western blot analysis [20].

### Immunofluorescence

Cells were harvested in a 1.5 ml Eppendorf tube, fixed with 3.7% paraformaldehyde for 10 minutes, and permeabilized with 0.25% Triton X-100 (Sigma, USA) for 15 minutes. Cells were placed on clean coverslips, followed by immunofluorescent staining using the indicated primary antibodies and respective fluorescent-labeled secondary antibodies (Alexa 488 and 594, Invitrogen, USA) as described earlier [20]. Fluorescent images were captured by a confocal microscope (model NLO710; Carl Zeiss, Germany).

### RNA isolation and RT-PCR

Total RNA was isolated using TRIzol reagent (Thermo Fisher Scientific, USA). Equal amounts of RNA (1 μg) were taken for cDNA synthesis using the iSCRIPT Bio-Rad cDNA synthesis kit (Bio-Rad, USA). Bio-Rad SYBR Green RT-PCR Master mix (Bio-Rad, USA) was used for real-time PCR. Actin was used as the internal control. The sequence of primers used is listed in Supplementary Table 5. The quantification of real-time data was done using the 2^-ΔΔCT^ method.

### Ubiquitination assay

HEK293T cells were cultured in a 60 mm dish and transfected at 50-60% confluency with 2 μg of each plasmid (Myc-WWP1 and GFP-RUNX1) with different ubiquitin linkage plasmid constructs (HA-WT Ub, HA-K0, HA-K6, HA-K11, HA-K27, HA-K29, HA-K33, HA-K48, HA-K63) using Lipofectamine 3000. After 12 h post-transfection, cells were replaced with complete cell culture medium for an additional 24 h before being harvested for immunoprecipitation analysis with the indicated antibodies.

### Cycloheximide pulse-chase assay

To assess protein stability, an equal number of WWP1-depleted K562 and KCL22 cells, with plasmids encoding WWP1 and RUNX1, were transfected into HEK293T cells. The cells were seeded for 24 h, and the indicated groups were treated with cycloheximide (100 μg/ml) followed by harvesting at various time points for western blot analysis.

### Apoptosis assay and DAD-FITC/PI staining

Apoptosis was assessed using the Dynamic Apoptosis Detection (DAD) kit (Cat#UR30120), UR Advanced Therapeutics, India, per the manufacturer’s instructions. K562 ShCtrl, ShWWP1, and WT cells (3x10^5^) were treated with Metformin (20 mM for 24 h) or compound C (1 μM for 6 h). Cells were incubated with 5 μl DAD reagent and 1 mg/ml propidium iodide (PI) solution in 800 μl PBS for 30 minutes at room temperature in the dark. Data acquisition was performed using a BD LSRFortesa flow cytometer, and analysis was conducted with FlowZo software [20].

### Synchronization of cells

To synchronize cells in the G2/M phase, K562 Sh ctrl and shWWP1 cells (5x10^5^ cells at 40% confluence) were treated with 50 μg/mL thymidine for 16 h. Cells were washed 3 times with 1X PBS, incubated in fresh RPMI-1640 medium for 12 h, and subsequently treated with nocodazole (100 ng/ml) for 16 h. Post-treatment, cells were released in pre-warmed RPMI-1640, and samples were collected at intervals up to 24 h. Synchronization was verified by flow cytometry.

### Cell cycle analysis

Cells were washed with ice-cold PBS containing 1% FBS, fixed with chilled 70% ethanol, and incubated at -20 °C for 30 minutes. Following three washes with ice-cold PBS, cells were resuspended in 200 μl of PBS with 100 μg of RNase A (Sigma-Aldrich) at 37 ^0^C for 30 minutes, then stained with 20 μl of propidium iodide (1 mg/ml) (Sigma–Aldrich) in 800 μl of PBS and incubated at 4 °C in the dark for 20 minutes. DNA content was analyzed using a FACS Aria (Becton Dickinson), and data were processed with CytoFlex software.

### Soft agar colony formation assay

A sterile base layer of 1.2% high-melting agar, mixed in a 1:1 ratio with RPMI-1640 (2X) and 20% FBS, was poured into six-well plates and allowed to solidify. A top layer of 0.6% low-melting agar mixed similarly with medium containing 1x10^5^ cells/well was added. Plates were incubated at 37 °C in 5% CO_2_ for 15-18 days, with fresh media added every 3 days. Colonies were fixed with 4% paraformaldehyde, stained with 0.05% crystal violet for 20 minutes, and washed to remove excess stain before being photographed.

### Site-directed mutagenesis

Site-directed mutations were introduced by PCR amplification using primers listed in Supplementary Table 2. The KLD (Kinase, Ligase, DpnI) enzyme reaction mix was used for post-PCR treatment per the kit protocol. The product was transformed into DH5α cells and plated on ampicillin-containing agar plates. Colonies were screened via colony PCR and confirmed with appropriate controls.

### Bacterial expression and protein purification

WWP1-WT, WWP1-T692A, and WWP1-T692D constructs were cloned into GST-tagged pGEX4T1 vectors. Recombinant proteins were expressed in BL21 *E. coli* cells induced with 1 mM IPTG at 37 °C overnight. Cell pellets were lysed by sonication in Sarcosyl, and GST-tagged proteins were purified using glutathione beads, eluting with 25 mM reduced glutathione. Eluted fractions were analysed by 10% SDS-PAGE.

### In vitro AMPK kinase assay

Kinase reactions were set up with 100 ng purified AMPK (EMD-Millipore, USA; Cat #14-840), 100 μM ATP, 1 mM dithiothreitol, 10 mM magnesium acetate, containing AMPK kinase buffer (20 mM HEPES (pH 7.4)), 1-5 μg of purified WWP1-WT, WWP1-T692A, and WWP1-T692D. Proteins were added to the reaction mixture, which was incubated at 37 °C for 30 minutes before being subjected to SDS-PAGE and western blotting.

### Tumor xenograft assay

All animal procedures were conducted in compliance with the ethical standards set by the Institutional Animal Ethics Review Committee (IAEC) and received approval (UH/IAEC/BMR/2021-1/04/R1). Six- to seven-week-old FOX N1 nude mice (n=6), equal mix of male and female, were obtained from the animal resource facility and were acclimated in individually ventilated cages (IVCs) for a week under specific pathogen-free conditions. Tumor xenografts were established by subcutaneous injection of 100 μl of PBS with 50% Matrigel (V/V) containing 1x10^7^ cells (Ctrl, WWP1 KD, RUNX1 KD, WWP1 overexpression, and metformin group). After six weeks, the animals were sacrificed, and the tumors were collected for further studies.

### Hematoxylin and eosin (H&E) staining

Formalin-fixed, paraffin-embedded (FFPE) tissues were subjected to standard sectioning. Hematoxylin and eosin staining. Stained sections were visualized using a Leica trinocular microscope.

### Statistical analysis

All the data were presented as mean ± SEM from at least three independent biological experiments (n=3) for each in vitro study. Statistical analyses were performed using one-way ANOVA, two-way ANOVA, or Student t-test, as appropriate. For the in vivo studies, each mouse represented a biological replicate. For the patient sample studies, each patient was considered as a biological replicate. A p-value < 0.05 was considered to be statistically significant. All analyses were performed using GraphPad Prism 8.0.

## Results

### High expression of WWP1 and RUNX1 correlates with poor prognosis of leukemic patients

WWP1 and RUNX1 have independently been implicated in the regulation of cellular proliferation, differentiation, and survival; however, their expression patterns, clinical relevance, and potential functional interplay in leukemic malignancies remain poorly defined. To establish the relevance of WWP1 and RUNX1 in leukemia, we first examined their expression profiles across leukemic cell lines and patient samples. For this, we analysed RNA-seq data from the Cancer Cell Line Encyclopedia (CCLE) database for cell lines from Acute myeloid leukemia (AML), Acute lymphoblastic leukemia (ALL), Chronic myeloid leukemia (CML), and Chronic lymphoblastic leukemia (CLL). We observed elevated transcript levels of both *WWP1* and *RUNX1* in ALL and CML cell lines compared to other cell lines (Figure 1A). Based on this observation, we further examined WWP1 and RUNX1 mRNA and protein expression in B-cells, HL-60, THP-1, K562, and Jurkat J6 cells. The mRNA expression of both genes was notably higher in K562 and Jurkat J6 cells (Figure 1B), with significantly higher protein levels in K562 cells than in B cells (Figure 1C and Supplementary Figure 1A). To investigate clinical significance, we assessed WWP1 and RUNX1 at the protein and mRNA levels in primary CML samples. Peripheral blood mononuclear cells (PBMCs) from healthy donors (n=2) and six primary CML patients (n=6) were studied. Despite a small sample size, both WWP1 and RUNX1 revealed differential expression in CML patient samples compared to healthy PBMCs (Figure 1D), confirming the clinical relevance of our findings. Then we analysed publicly available TCGA datasets using the Gene Expression Profiling Interactive Analysis (GEPIA) web server. Both *WWP1* and *RUNX1* were significantly upregulated in AML patient samples (n=173) compared with normal controls (n=70) (Supplementary Figure 1B). Notably, correlation analysis revealed a strong positive correlation between *WWP1* and *RUNX1* expression (R=0.82, p<0.0001), suggesting a potential regulatory or functional interaction in leukemia progression (Supplementary Figure 1C). Kaplan-Meier survival analysis of AML patients stratified by high (n=53) versus low (n=53) *WWP1* transcript levels and high (n=76) versus low (n=76) *RUNX1* transcript levels (TCGA data sets) results in poor overall survival of AML patients (P=0.15 for *WWP1* and 0.032 for *RUNX1*; Supplementary Figure 1D & E). Collectively, these results demonstrate that WWP1 and RUNX1 are highly expressed in leukemic settings and may represent a functionally relevant oncogenic axis.

**Figure 1:**
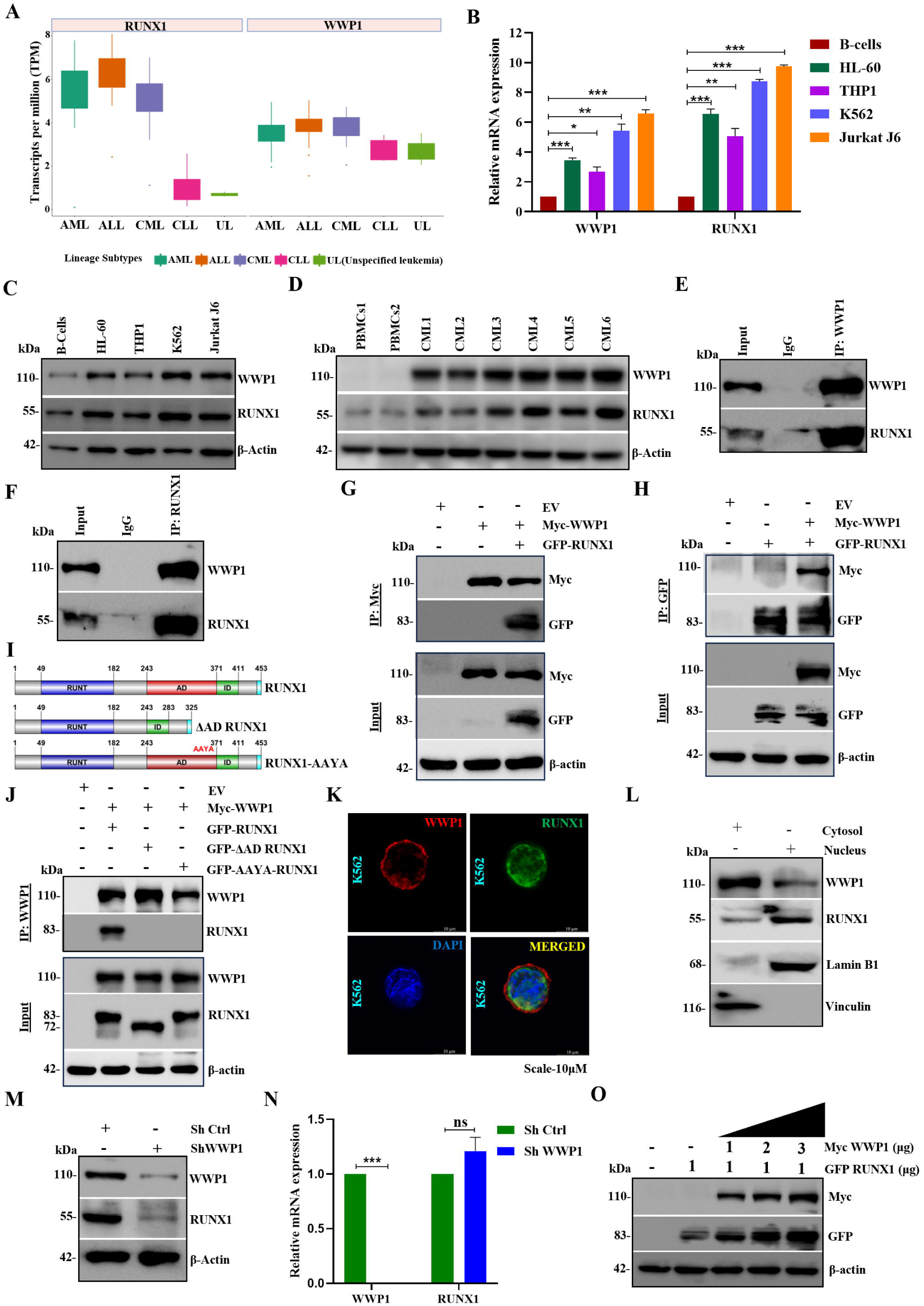
Elevated expression and interaction of WWP1 and RUNX1 in CML. **A.** Gene expression profiles of *WWP1* and *RUNX1* across various human leukemic cell lines were obtained from the CCLE RNA-seq database (transcripts per million (TPM) were quantified using RSEM, followed by log2 transformation (pseudo-count = 1)). Box plots depict lineage-wise average expression, highlighting elevated levels in CML and ALL cell lines. **B.** Quantitative RT-PCR analysis of WWP1 and RUNX1 mRNA expression in leukemia cell lines relative to B-cells. **C.** Immunoblot analysis of endogenous WWP1 and RUNX1 protein expression in leukemia cell lines relative to B-cells **D.** Protein expression of WWP1 and RUNX1 in PBMCs from healthy donors and primary CML patient cells. **E & F.** Endogenous interaction between WWP1 and RUNX1 in K562 cells was demonstrated by co-immunoprecipitation using anti-WWP1 (E) and anti-RUNX1 (F) antibodies, followed by immunoblotting with the indicated antibodies. **G** & **H.** Interaction between ectopically expressed Myc-WWP1 and GFP-RUNX1 in HEK293T cells confirmed by reciprocal co-immunoprecipitation using anti-Myc (G) and anti-GFP (H) antibodies. **I.** Schematic representation of full-length RUNX1, the ΔAD deletion mutant, and the AAYA mutant used for interaction mapping. **J**. Co-immunoprecipitation analysis of the indicated RUNX1 constructs, identifying the PPPY as the major region required for interaction with WWP1. **K.** Immunofluorescence analysis showing the subcellular co-localization of endogenous WWP1 (Alexa-546) and RUNX1 (Alexa-488) in K562 cells. Nuclei were counterstained with DAPI. Scale bar, 10 μm. **L**. Cytoplasmic and nuclear fractionation of K562 cells demonstrating the subcellular distribution of WWP1 and RUNX1. **M.** Immunoblot analysis of WWP1 and RUNX1 expression following shRNA-mediated knockdown of WWP1. **N.** RT-PCR analysis showing the effect of WWP1 knockdown on RUNX1 mRNA expression. **O.** Dose-dependent overexpression of Myc-WWP1 in K562 cells and its effect on endogenous RUNX1 protein expression. Data are presented as mean ± SEM from three independent biological experiments (n=3). In panel B, the relative mRNA expression of WWP1 and RUNX1 in each leukemia cell line was individually compared with B-cells. In panel N, RUNX1 mRNA expression in shWWP1 cells was compared with the corresponding shCtrl cells. Statistical significance was determined using an unpaired, two-tailed Student’s t-test. Statistical significance is indicated as *p<0.05, **p<0.01, ***p<0.001; ns: not significant.

### WWP1 interacts with RUNX1 and regulates it at post-translational levels

Previous studies have reported that RUNX1 may interact with nine E3 ubiquitin ligases, including WWP1 [13]. WWP1 is known to target substrates with PPxY-containing motifs [25, 26]. To investigate whether RUNX1 contains such a motif, we analyzed the protein sequence using NCBI resources and confirmed the presence of a PPxY motif in RUNX1 (Supplementary Figure 2A). To experimentally validate the interaction between WWP1 and RUNX1, we performed a co-immunoprecipitation assay in HEK293T cells, which have low or negligible expression of both proteins. We transiently co-expressed Myc-WWP1 and GFP-RUNX1 and immunoprecipitated using both anti-Myc and anti-GFP antibodies, confirming a strong interaction between the two proteins (Figure 1G&H). We further validated this interaction at endogenous levels in K562 and KCL22 cells using anti-WWP1 and anti-RUNX1 antibodies (Figure 1E&F, and Supplementary Figure 2B&C). Similarly, endogenous co-immunoprecipitation assays in HL-60 and Jurkat J6 cell lines also showed interaction between the two proteins (Supplementary Figure 2D-G).

We next asked which RUNX1 domain mediates its interaction with WWP1. Earlier reports have shown that WWP1 interacts with RUNX2 [25]. We analysed the structural domains of WWP1 and RUNX1 using UniProt (Supplementary Figure 2H&I). Protein-protein docking using the HDOCK server predicted a strong interaction between the WW3 domain of WWP1 and the PPPY motif of RUNX1 (Supplementary Figure 2J). To experimentally validate this prediction, we generated AD domain-deleted (ΔAD) and AAYA mutant constructs of RUNX1 using a site-directed mutagenesis approach (Figure 1I). Co-transfection experiments in HEK293T cells demonstrated that only wild-type RUNX1 interacted with WWP1, whereas the ΔAD and AAYA mutants failed to do so (Figure 1J), confirming the domain-specific interaction.

To investigate subcellular localization, we performed immunofluorescence staining and observed that both WWP1 and RUNX1 are colocalized in the cytoplasm as well as in the nucleus, with WWP1 enriched in the cytoplasm and RUNX1 in the nucleus (Figure 1K). These findings were further corroborated by nuclear-cytoplasmic fractionation experiments (Figure 1L). As WWP1 functions as an E3 ubiquitin ligase and is positively regulated with RUNX1 in leukemia patients (Supplementary Figure 1C), we next examined whether it regulates RUNX1 at the transcriptional or post-translational level. We generated stable K562 and KCL22 cell lines with lentiviral-mediated shRNA knockdown of WWP1 (Supplementary Figure 2K&L). Knockdown of WWP1 led to a marked reduction in RUNX1 protein levels (Figure 1M and Supplementary Figure 2L&M) without any significant change in RUNX1 mRNA levels (Figure 1N), suggesting post-translational regulation. To further validate this, we have transiently transfected HEK293T cells with increasing amounts of WWP1 along with RUNX1-expressing plasmids. WWP1 overexpression led to a dose-dependent increase in RUNX1 protein levels (Figure 1O & Supplementary Figure 2N), again without affecting RUNX1 mRNA levels (Supplementary Figure 2O). Taken together, these data demonstrate that WWP1 physically interacts with the PPPY motif of RUNX1, and regulates RUNX1 protein stability at the post-translational levels.

### WWP1 promotes K63-linked polyubiquitination and stabilizes RUNX1

To explore the molecular relationship between WWP1 and RUNX1 in CML, we first confirmed a physical interaction between the two proteins. In silico analysis using UbiBrowser 2.0 predicted RUNX1 as a potential WWP1 substrate, with a high confidence score (0.916), supporting the likelihood of WWP1-mediated RUNX1 ubiquitination (Figure 2A and Supplementary Figure 3A). To experimentally validate this, we evaluated total cellular ubiquitination levels in K562 and KCL22 cells following WWP1 knockdown using either shRNA or pharmacological inhibition with Indole-3-Carbinol (I3C), a known WWP1 inhibitor. Both interventions led to a marked reduction in overall ubiquitination of RUNX1 (Figure 2B and Supplementary Figure 3B&C). Similar results were observed upon WWP1 knockdown or inhibition with I3C in Jurkat J6 and HL-60 leukemia cell lines, indicating the conserved role of WWP1-RUNX1 ubiquitination across hematological malignancies (Supplementary Figure 3D-G).

**Figure 2:**
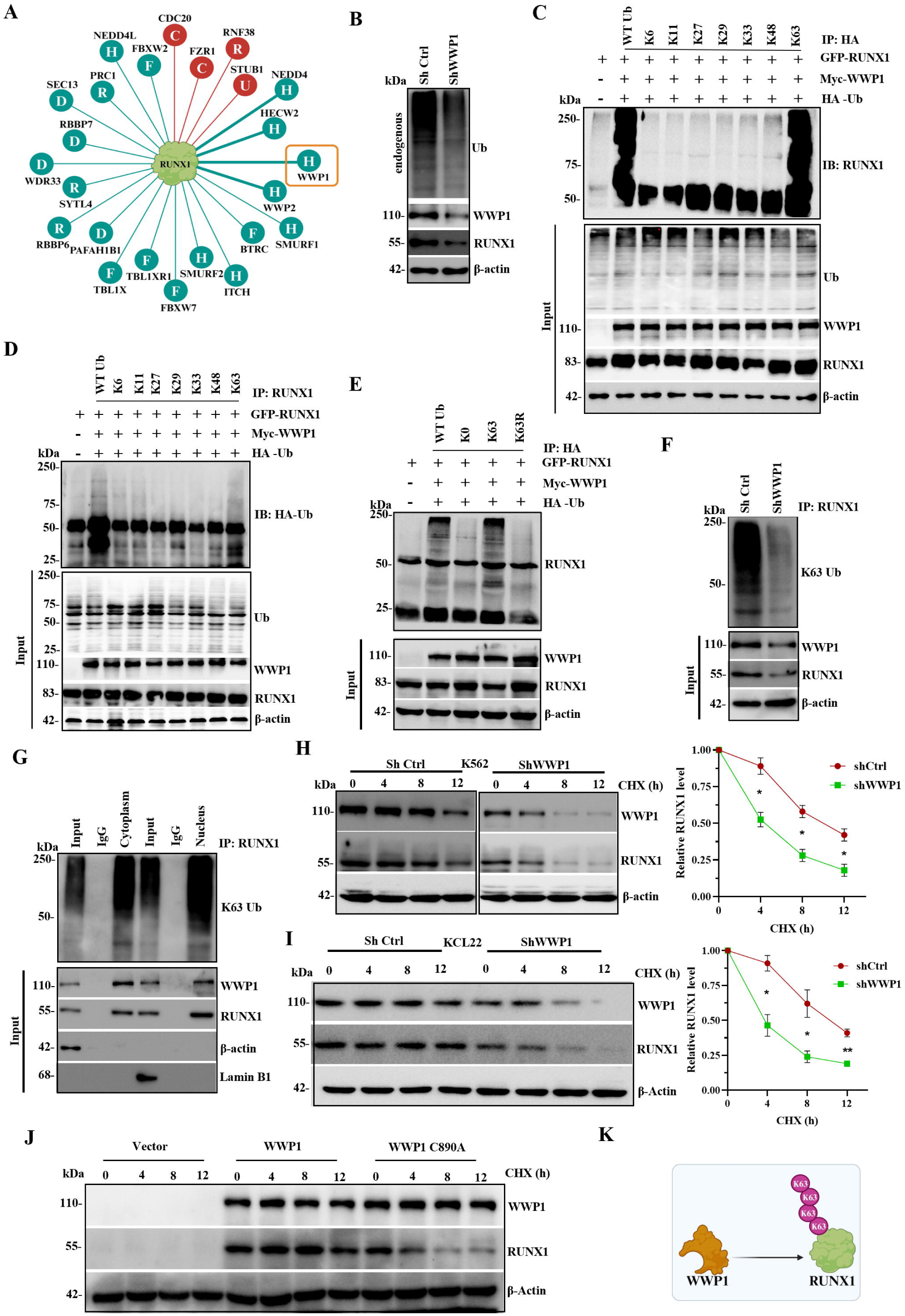
WWP1 promotes K63-linked polyubiquitination and stabilizes RUNX1 protein. **A.** UbiBrowser 2.0 prediction identifying WWP1 as a candidate E3 ubiquitin ligase for RUNX1. **B.** Immunoblot analysis of total ubiquitinated proteins and the expression of WWP1 and RUNX1 in shCtrl and shWWP1 K562 cells. **C & D.** HEK293T cells were co-transfected with the indicated plasmids and the indicated ubiquitin lysine mutants. RUNX1 ubiquitination was analyzed by immunoprecipitation with anti-HA (C) and anti-RUNX1 (D) followed by immunoblotting using the indicated antibodies. **E.** HEK293T cells were co-transfected with the indicated plasmids and the indicated ubiquitin lysine mutants. RUNX1 ubiquitination was analyzed by immunoprecipitation using anti-HA antibodies. **F.** Endogenous K63-linked ubiquitination of RUNX1 was examined in shCtrl and shWWP1 K562 cells by immunoprecipitation with anti-RUNX1, followed by immunoblotting using a K63 linkage-specific ubiquitin antibody. **G.** Subcellular localization of K63-linked ubiquitinated RUNX1 was analyzed by immunoprecipitation of cytoplasmic and nuclear fractions using anti-RUNX1 antibodies. **H** & **I.** CHX chase assay showing the stability of endogenous RUNX1 protein in shCtrl and shWWP1 K562 cells (H) and KCL22 cells (I). Right panel, densitometric quantification of RUNX1 protein levels normalized to -actin. The RUNX1-to-actin ratio at the initial (0 h) time point was defined as 1, and relative changes in RUNX1 levels at subsequent time points were calculated with reference to this baseline. **J.** HEK293T cells were transfected with empty vector, Myc-WWP1, or the catalytically inactive WWP1 C890A mutant, followed by CHX treatment to assess the effect of WWP1 E3 ligase activity on RUNX1 protein stability. **K.** Proposed model illustrating WWP1-mediated K63-linked polyubiquitination and stabilization of RUNX1. Data are presented as mean ± SEM from three independent biological experiments (n=3). RUNX1 protein levels at each CHX treatment time point in shWWP1 cells were compared with the corresponding shCtrl cells using two-way ANOVA followed by Sidak’s multiple comparisons test. Statistical significance is indicated as *p<0.05, **p<0.01; ns: not significant.

Protein degradation is generally marked by K48-linked ubiquitination [13], whereas stabilization and transcriptional regulation are mediated by K63 [11, 16]. To characterize the specific type of ubiquitination involved, we conducted ubiquitination assays in HEK293T cells co-expressing WWP1, RUNX1, and HA-tagged ubiquitin mutants where all lysines except one were substituted with arginine. Pull-down using anti-HA and anti-RUNX1 antibodies demonstrated that WWP1 specifically promotes K63-linked polyubiquitination of RUNX1 (Figure 2C-D). This was confirmed using K63-only, K63R, and lysine-null (K0) mutant ubiquitin constructs-only the K63 construct supported WWP1-mediated RUNX1 ubiquitination (Figure 2E). Since WWP1 functions via its catalytically active HECT domain, we tested a ligase-inactive mutant (C890A). Unlike wild-type (WT) WWP1, the C890A mutant failed to promote K63-linked ubiquitination of RUNX1 (Supplementary Figure 3H), establishing the requirement of WWP1’s ligase activity for this modification. Endogenous RUNX1 immunoprecipitation followed by probing for K63-linked ubiquitin confirmed that WWP1 depletion significantly reduced RUNX1 ubiquitination (Figure 2F). Subcellular fractionation revealed that ubiquitinated RUNX1 predominantly localised to the nuclear compartment (Figure 2G), suggesting a potential role in transcriptional regulation rather than proteasomal degradation.

Next, to determine whether WWP1-mediated K63 ubiquitination influences RUNX1 protein stability, we performed a cycloheximide (CHX) chase assay in K562 and KCL22 cells. In control cells, RUNX1 turnover proceeded slowly, whereas WWP1 knockdown led to a substantial (∼6-fold) increase in RUNX1 turnover (Figure 2H–I). Conversely, ectopic co-expression of WWP1 and RUNX1 decreases protein turnover of RUNX1 in HEK293T cells over time with CHX treatment, but in the presence of the WWP1 C890A mutant, RUNX1 protein turnover increases over time (Figure 2J and Supplementary Figure 3I). Overall, these data show that WWP1 promotes K63-linked polyubiquitination of RUNX1, which increases RUNX1 protein stability rather than targeting it for proteasomal degradation. This non-canonical ubiquitination likely promotes RUNX1’s nuclear retention and functional lifespan (Figure 2K), providing insight into the mechanism of RUNX1 regulation in leukemogenesis.

### WWP1 Silencing reduces CML cell growth by causing G2/M arrest and apoptosis

To determine the functional relevance of WWP1 in CML, we generated stable WWP1-knockdown cell lines and assessed their proliferative potential. WWP1 loss resulted in a considerable reduction in cell proliferation relative to control cells (Figure 3A). To confirm these findings, we used clonogenic assays, which indicated a significant decrease in both colony size and number in WWP1-silenced cells (Figure 3B&C and Supplementary Figure 4A&B), highlighting WWP1’s crucial role in maintaining CML oncogenic potential.

**Figure 3:**
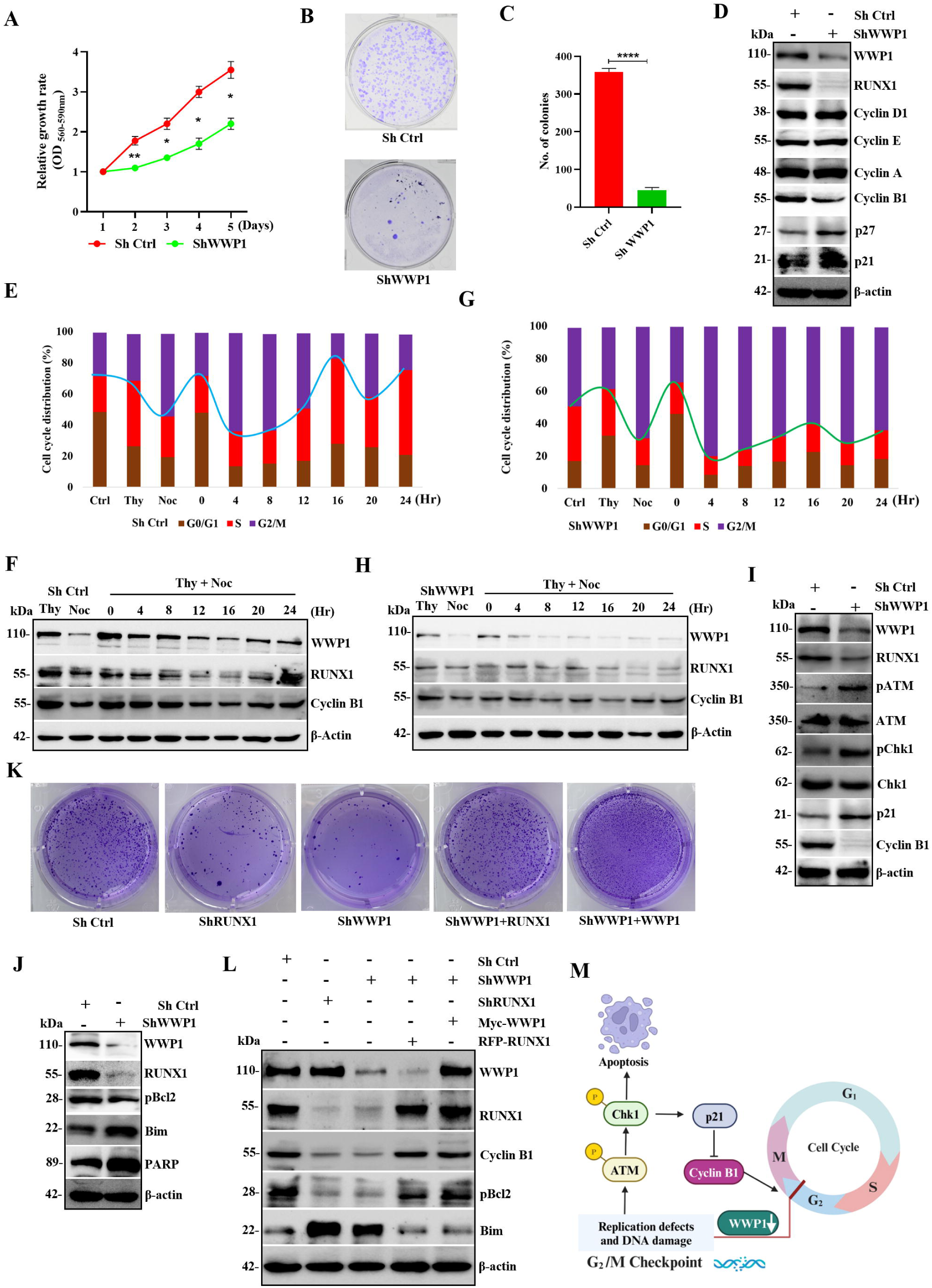
WWP1 promotes leukemic cell proliferation by regulating G2/M cell-cycle progression and cell survival. **A.** Cell proliferation of shCtrl and shWWP1 K562 cells was determined over five consecutive days using the PrestoBlue assay. **B.** Representative images from colony formation by shCtrl and shWWP1 K562 cells. **C.** Quantification of colonies formed by shCtrl and shWWP1 K562 cells. **D.** Immunoblot analysis of the indicated protein expression following WWP1 knockdown in K562 cells. **E & G.** Cell cycle distribution of synchronized shCtrl (E) and shWWP1 (G) K562 cells following thymidine-nocodazole block and release at the indicated time points. **F & H.** Immunoblot analysis of WWP1, RUNX1 and Cyclin B1 protein expression during cell cycle progression in synchronized shCtrl (F) and shWWP1 (H) K562 cells. **I.** Immunoblot analysis of DNA damage check point proteins following WWP1 knockdown. **J.** Immunoblot analysis of apoptosis-associated proteins in shCtrl and shWWP1 K562 cells. **K.** Representative colony formation assay demonstrating the effects of WWP1 knockdown, RUNX1 knockdown, WWP1 overexpression, and rescue experiments on clonogenic growth. **L.** Immunoblot analysis of WWP1, RUNX1, Cyclin B1, pBcl2, and Bim expression in WWP1 knockdown, RUNX1 knockdown, WWP1 overexpression, and rescue conditions. **M.** Schematic model illustrating that WWP1-mediated stabilization of RUNX1 promotes G2/M cell-cycle progression and supresses apoptosis, whereas WWP1 depletion activates the ATM-Chk1-p21 signaling pathway. Leading to G2/M checkpoint activation and apoptotic cell death. Data are presented as mean ± SEM from three independent biological experiments (n=3). (A) Cell proliferation of shWWP1 cells was compared with the corresponding shCtrl cells at each time point using a two-way ANOVA followed by Sidak’s multiple comparisons test. (C) Colony numbers in shWWP1 cells were compared with the corresponding shCtrl cells using an unpaired, two-tailed Student’s t-test. Statistical significance is indicated as *p<0.05, **p<0.01, ****p<0.0001.

We subsequently examined the role of WWP1 in cell proliferation by analysing cell-cycle checkpoint markers using western blotting. Our results showed reduced cyclin B1 levels in WWP1-knockdown cells, a G2/M phase-specific cyclin, compared to controls (Figure 3D and Supplementary Figure 4C). To further confirm the Western blot results, we synchronized the K562 cells by treating them with double thymidine-nocodazole and releasing them as described in the methods. Cells were collected periodically at different time points and analyzed. Our results show that WWP1 knockdown cells took 24 h to pass through the G2/M phase, whereas controls passed within 16 h (Figure 3E &F and Supplementary Figure 5A). Thereby underscoring the essential function of WWP1 in the G2/M transition.

Since the G2/M checkpoint is critical in response to DNA damage and triggers apoptosis, we subsequently assessed the activation of the DNA damage response signaling proteins ATM and CHK1 in WWP1-knockdown cells compared to controls (Figure 3I and Supplementary Figure 4C), indicating activation of the DNA damage response signaling. We then measured apoptosis using the DAD(R) apoptosis detection kit and confirmed that 50.74% of WWP1 knockdown cells exhibited apoptosis compared to controls (6.27%). Knocking down WWP1 cells showed reduced anti-apoptotic markers and increased pro-apoptotic marker expression (Figure 3J and Supplementary Figure 4D&E). In summary, these findings reveal that WWP1 promotes CML growth by altering the G2/M cell cycle and inhibiting apoptosis.

### RUNX1 is required for the oncogenic function of WWP1 in CML

WWP1 is a well-known oncogene in a number of solid tumors, and RUNX1 has been implicated as an oncogenic driver in blood cancers. To determine the necessity of RUNX1 for the oncogenic activity of WWP1 in CML, we performed a series of in vitro functional tests utilising K562 and KCL22 cells subjected to either WWP1 or RUNX1 knockdown. PrestoBlue cell viability assays demonstrated that knockdown of WWP1 and RUNX1 significantly reduced cell proliferation relative to the control. Ectopic overexpression of WWP1 in WWP1-depleted cells and RUNX1 in WWP1 knockdown cells partially restored proliferative ability compared to respective experimental controls (Supplementary Figure 4F&G). To elucidate the role of RUNX1 as a downstream effector of WWP1 in promoting leukemic proliferation, we observed a significant increase in colonies in a colony-forming assay following ectopic overexpression of WWP1 and RUNX1 in WWP1-knockdown K562 and KCL22 cells (Figure 3K and Supplementary Figure 4H-J). The data show that RUNX1 is necessary for WWP1 to cause CML development.

To clarify the mechanism driving this proliferative effect, we evaluated the expression of apoptotic regulators. Western blot analyses revealed elevated levels of pro-apoptotic proteins and reduced levels of anti-apoptotic markers in WWP1 and RUNX1-knockdown K562 and KCL22 cells. Conversely, cells overexpressing both WWP1 and RUNX1 exhibited increased anti-apoptotic signaling and reduced expression of pro-apoptotic proteins in both K562 and KCL22 cells (Figure 3L and Supplementary Figure 4K&L). Collectively, these findings demonstrate that RUNX1 is a critical effector of WWP1-mediated oncogenic signaling in CML and that WWP1 promotes leukemic cell proliferation and survival, at least in part, by regulating apoptosis and DNA damage response pathways (Figure 3M).

### WWP1 is a direct substrate of AMPK and undergoes phosphorylation at Thr692 position

WWP1 promotes the growth and progression of several solid tumors, including AML, and we have demonstrated its role in CML by promoting proliferation through ubiquitination of RUNX1. Our recent findings showed that p-RUNX1 inhibits CML cell proliferation [20], prompting us to explore upstream regulators of WWP1 that might modulate this axis. Given the role of AMPK as a central energy sensor and regulator of metabolic stress responses [27], we hypothesized that AMPK may regulate WWP1 activity. To test this, we first performed *in silico* predictions using the PhosphoMotif Finder tool, which identified a high-confidence AMPK substrate motif within amino acids 687–696 of WWP1, with threonine 692 as the predicted phosphorylation site (Figure 4A and Figure 5A). The motif contains a preferred basic residue at the –3 position and hydrophobic residues (leucine at –5 and methionine at +4), which are characteristic of AMPK substrates [24, 27, 28] and have been validated earlier by site-directed mutagenesis and molecular modeling studies. This prediction was further supported by Net-Phos 3.1, which yielded a high phosphorylation score for Thr692 (Score-0.854). Importantly, other predicted kinases targeting this motif lacked consensus homology or conservation, whereas the AMPK recognition motif in WWP1 is conserved across known AMPK substrates and species (Figure 4B&4C).

**Figure 4:**
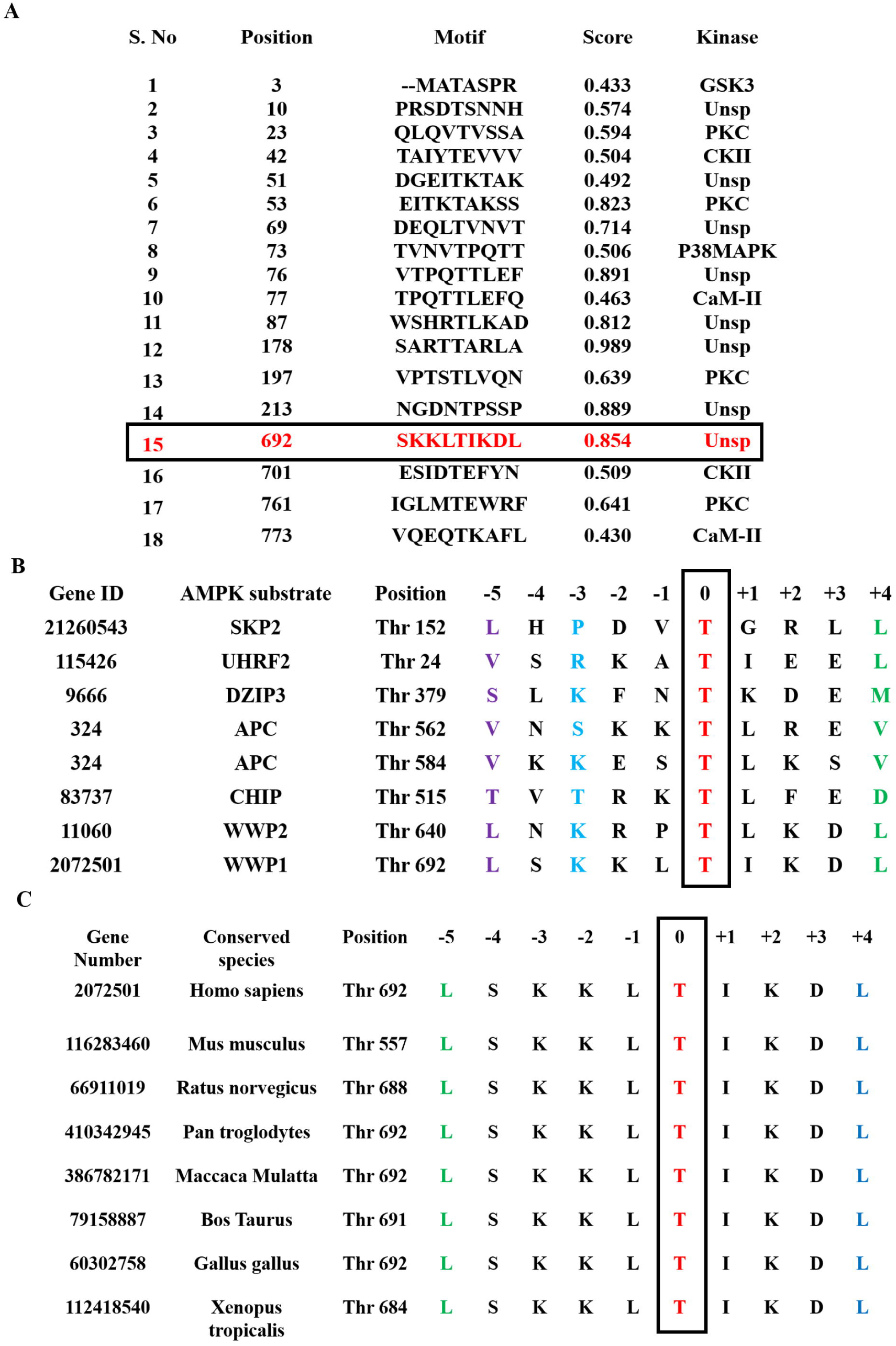
In silico prediction identifies Thr692 as a putative AMPK phosphorylation site in WWP1. **A.** Prediction of potential phosphorylation sites in WWP1 using the PhosphoMotif Finder tool identified a high-confidence AMPK consensus motif spanning amino acids 687-696, with Thr692 predicated as the putative AMPK phosphorylation residue (highlighted in red). **B**. Sequence alignment of the predicted WWP1 AMPK phosphorylation motif with established AMPK substrates using NetPhos 3.1 demonstrates conservation of the consensus AMPK recognition sequence, supporting Thr692 as a potential phosphorylation site. **C**. Cross-species multiple sequence alignment of the WWP1 region encompassing Thr692 using ClustalW2 reveals evolutionary conservation of the predicted AMPK phosphorylation motif among different vertebrate species.

**Figure 5:**
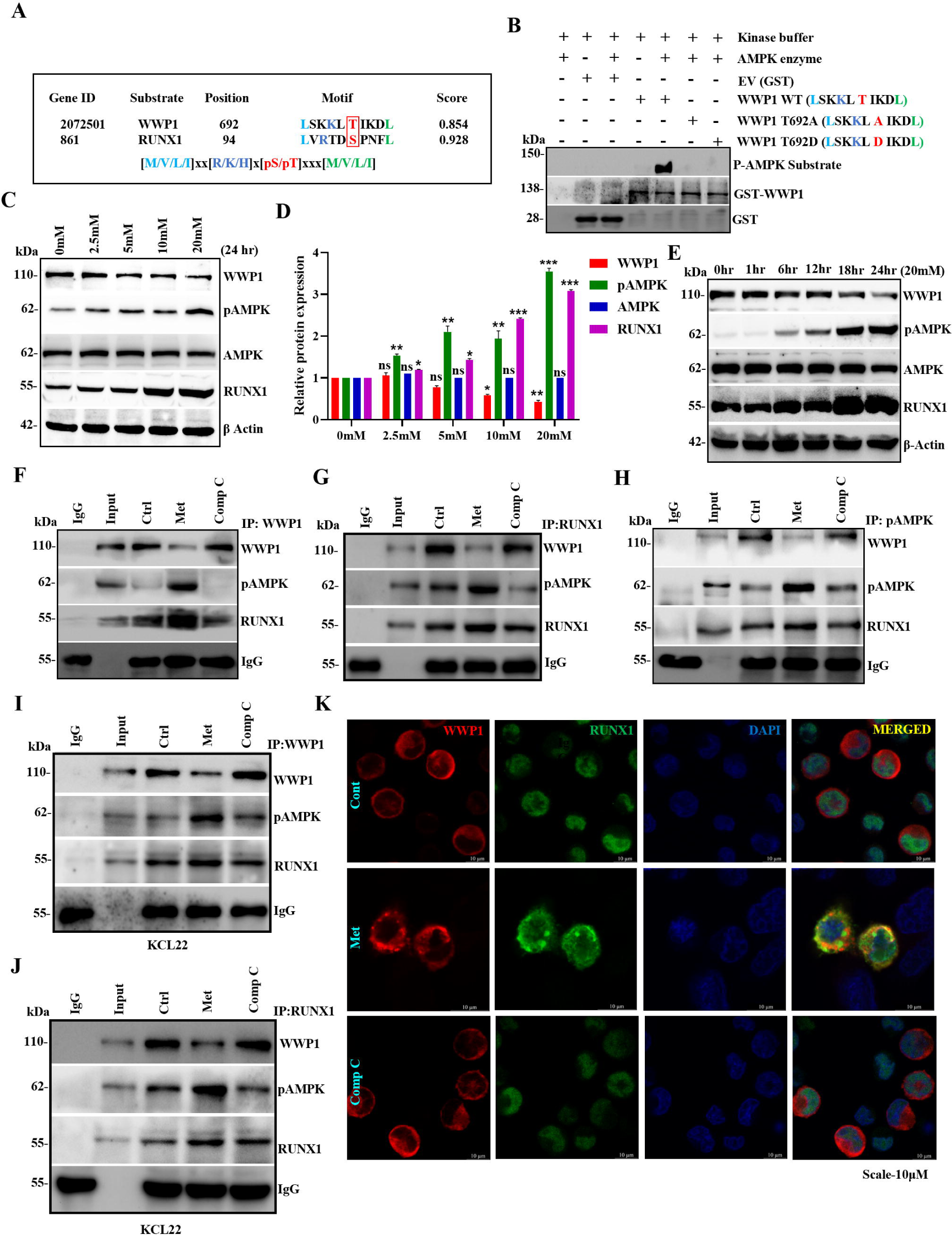
AMPK phosphorylates WWP1 at Thr692 and negatively regulates WWP1-mediated RUNX1 signaling. **A.** Prediction of putative AMPK phosphorylation sites within WWP1 using PhosphoMotif finder. Thr692 was identified as the conserved AMPK phosphorylation site. **B.** *In vitro* kinase assay followed by immunoblotting with phospho-AMPK substrate-specific antibody using partially purified bacterial-expressed WWP1 and its domain-specific mutants. **C& D.** Immunoblot analysis of indicated protein expression in K562 cells treated with increasing concentrations of metformin for 24 h (C) and the densitometric quantification of the indicated protein expression following metformin treatment. Protein levels were normalized to -actin and expressed relative to the untreated control (D). **E**. Immunoblot analysis of indicated protein expression following treatment with metformin at different time points. **F.** Immunoprecipitation of endogenous WWP1, **G.** RUNX1, **H.** p-AMPK from K562 cells treated with metformin (20 mM) for 24 h and compound C (5 μM) for 6 h, followed by immunoblotting with the indicated antibodies. **I.** Immunoprecipitation of endogenous WWP1 and **J.** RUNX1 in KCL22 cells treated with metformin (20 mM) or compound C (5 μM) for 6 h, followed by immunoblotting with the indicated antibodies. **K.** Immunofluorescence analysis for assessing the subcellular localization of endogenous WWP1 (Alexa-546) and RUNX1 (Alexa-488) proteins in K562 cells following treatment with metformin (20 mM) and Comp C (5 μM). Nuclei were counterstained with DAPI. Scale bar, 10 μm. Data are presented as mean ± SEM from three independent biological experiments (n=3). Protein expression levels of the indicated protein at each metformin concentration were individually compared with the corresponding untreated control using one-way ANOVA followed by Dunnett’s multiple comparison test. Statistical significance is indicated as *p<0.05, **p<0.01, ***p<0.001; ns: not significant.

To experimentally validate the *in silico* prediction, we performed an *in vitro* AMPK kinase assay using a cell-free system with wild-type (WT), phospho-null (T692A), and phospho-mimic (T692D) WWP1 proteins (Supplementary Figure 6A). Immunoblotting using an anti-phospho AMPK substrate motif antibody showed phosphorylation of only the WT-WWP1 protein, confirming Thr692 as the specific AMPK target site (Figure 5B). Consistently, treatment of K562 cells with metformin, a pharmacological AMPK activator [29, 30], led to a significant reduction in cell viability (Supplementary Figure 6B). Dose- and time-dependent treatment with metformin showed progressive downregulation of WWP1 protein levels and concurrent increase of RUNX1 levels in K562 and KCL22 cell lines (Figure 5C-E and Supplementary Figure 6C-E), while WWP1 mRNA levels remained unchanged, indicating post-translational regulation (Supplementary Figure 6F). Similar results were obtained with I3C, another known AMPK activator [31], reinforcing the inverse relationship between WWP1 and RUNX1 expression (Supplementary Figure 6G-I). To assess AMPK targeting of WWP1 phosphorylation in cells, probing with a pAMPK-substrate-specific antibody following metformin treatment increased AMPK-specific WWP1 phosphorylation (Supplementary Figure 6J). Co-immunoprecipitation analysis revealed that AMPK co-precipitated with both WWP1 and RUNX1 when treated with metformin, and this was disrupted by treatment with the AMPK inhibitor, compound C, in K562 and KCL2 cell lines (Figure 5F-J). Immunofluorescence imaging reveals that WWP1 is localized to the cytoplasm and RUNX1 in the nucleus. Upon treatment with metformin, WWP1 formed cytoplasmic puncta indicative of lysosomal targeting, which was reversed by compound C treatment (Figure 5K). These findings collectively demonstrate that WWP1 is a direct substrate of AMPK and is specifically phosphorylated at Thr692. AMPK-induced phosphorylation of WWP1 leads to post-translational regulation, thereby uncovering a novel regulatory axis, WWP1-RUNX1, in CML progression.

### AMPK-induced phosphorylation facilitates the lysosomal degradation of WWP1 and the stability of RUNX1 in CML

Prior results indicated that metformin treatment markedly decreased WWP1 protein levels, as demonstrated by immunoblotting, and elicited prominent cytoplasmic puncta in immunofluorescence examination, indicative of WWP1 degradation (Figure 5K). To further examine this process, we overexpressed RUNX1 in conjunction with either WT WWP1 or phospho-mutants (T692A and T692D) and then treated with metformin. Immunoblot examination demonstrated a notable decrease in WWP1 protein levels in both WT and phospho-mimetic T692D mutants, whereas no reduction was observed in the phospho-deficient T692A mutant (Figure 6A). RUNX1 levels increased in both the WT and T692D groups after metformin therapy, indicating that AMPK-mediated phosphorylation at Thr692 promotes WWP1 breakdown and increases RUNX1 stability. To assess RUNX1 stability more directly, we transiently overexpressed WWP1 and treated cells with metformin. In keeping with earlier findings, RUNX1 levels remained stable despite decreasing WWP1 levels after treatment (Supplementary Figure 7A&B), demonstrating that AMPK activation increases RUNX1 stability via inhibiting WWP1. This evidence supports our hypothesis that WWP1 promotes leukemogenic RUNX1 turnover, while AMPK activation inhibits this process, resulting in anti-leukemic effects.

**Figure 6:**
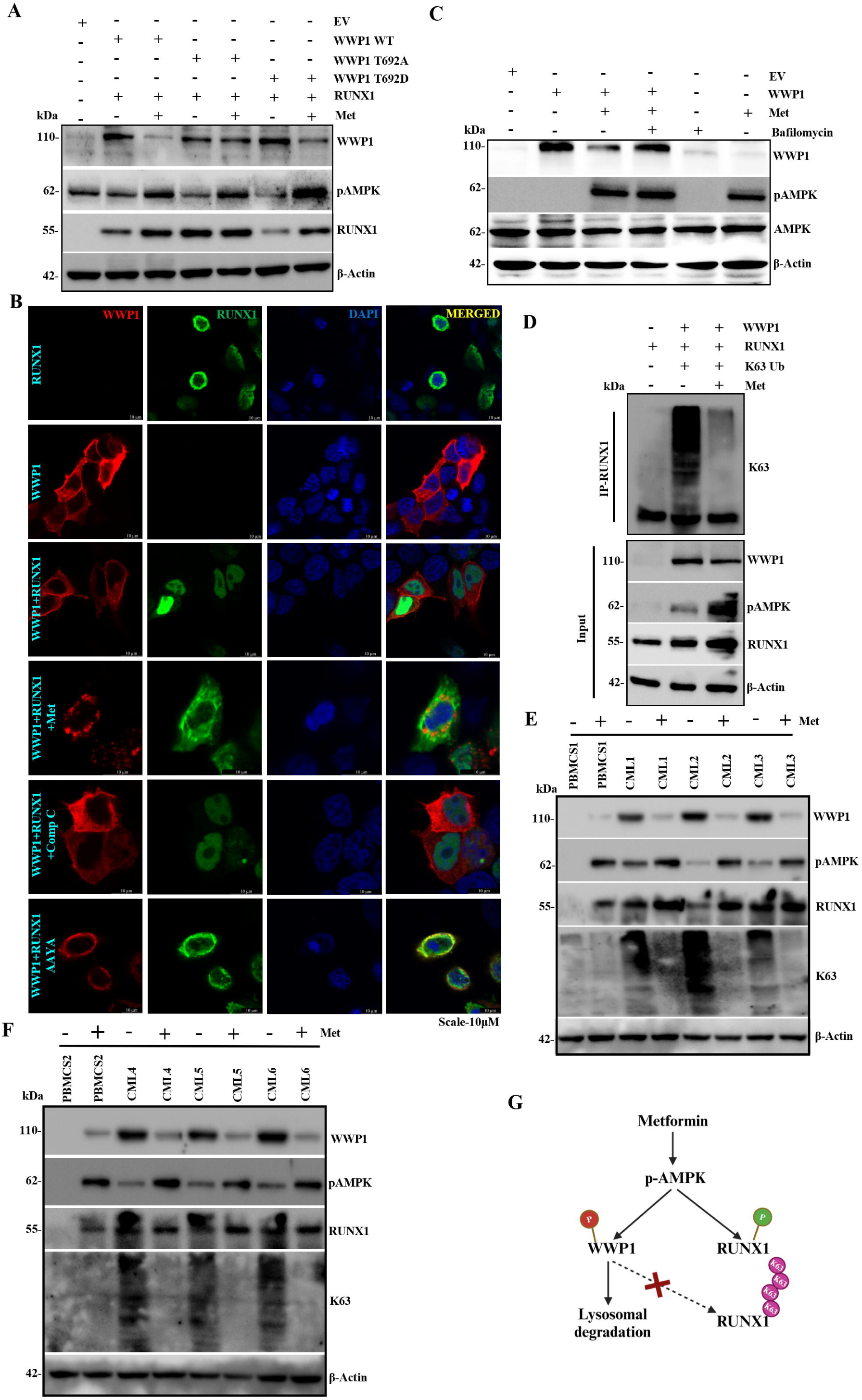
AMPK facilitates lysosomal degradation of WWP1 and stabilizes RUNX1 in CML. **A.** HEK293T cells were transfected with the indicated WWP1 constructs and treated with metformin (20 mM), followed by immunoblot analysis with the indicated antibodies. **B.** Immunofluorescence analysis showing the subcellular localization of WWP1 (Alexa-546) and RUNX1 (Alexa-488) in HEK293T cells expressing the indicated constructs following metformin (20 mM) and compound C (μM) treatment. Nuclei were counterstained with DAPI. Scale bar, 10 μm. **C.** HEK293T cells expressing the indicated WWP1 constructs were treated with metformin (20 mM) for 24 h in the presence or absence of bafilomycin A1 (5 μM) for 6 h. Whole-cell lysates were subjected to immunoblot analysis. **D.** HEK293T cells expressing the indicated plasmids were treated with metformin (20 mM). Denatured whole-cell lysates were immunoprecipitated with an anti-RUNX1 antibody, followed by immunoblot analysis using the indicated antibodies. **E &F.** Primary PBMCs with CML were treated with metformin (20 mM) for 24 h. Whole-cell lysates were subjected to immunoblot analysis using the indicated antibodies. **G.** Schematic model depicting AMPK activation promotes phosphorylation-dependent lysosomal degradation of WWP1, thereby reducing WWP1-mediated K63-linked ubiquitination of RUNX1.

We subsequently examined the subcellular localization of RUNX1 mimicking the ubiquitinated and phosphorylated conditions. The immunofluorescence analysis revealed that increased nuclear localization of RUNX1 (Ubiquitinated) compared to phosphorylated RUNX1 (non-ubiquitinated). Upon treatment with metformin, WWP1 undergoes cytoplasmic degradation, and RUNX1 is sequestered in the cytoplasm. Treatment of compound C restores WWP1 in the cytoplasm, and RUNX1 translocates to the nucleus in HEK293T cells (Figure 6B), confirming that AMPK activity governs these changes. To clarify, we overexpressed WWP1 in HEK293T cells and treated them with metformin, either independently or in combination with the proteasomal inhibitor MG132 or the lysosomal inhibitor bafilomycin A1. Notably, treatment with bafilomycin A1 significantly restored WWP1 levels, but not with MG132 (Figure 6C and Supplementary Figure 7C), suggesting that WWP1 is subjected to lysosomal rather than proteasomal degradation.

Immunoprecipitation of RUNX1 showed that metformin treatment decreased its polyubiquitination (Figure 6D), thereby reinforcing the hypothesis that AMPK activation interferes with WWP1-mediated RUNX1 ubiquitination. To enhance the therapeutic significance of our findings, primary CML patient cells were treated with metformin. Metformin treatment led to activation of AMPK, which diminished WWP1 expression, significantly increased RUNX1 phosphorylation, and reduced K63-linked ubiquitination (Figure 6E&F). These studies indicate that metformin efficiently regulates the WWP1-AMPK-RUNX1 signaling axis in primary CML patient cells, hence reinforcing the translational relevance of our mechanistic findings. Together, these findings establish a novel AMPK-WWP1-RUNX1 axis in which AMPK phosphorylates WWP1 at T692, thereby targeting it for lysosomal degradation (Figure 6G). This degradation diminishes RUNX1 ubiquitination, leading to its stabilization through phosphorylation and potential anti-leukemic effects in CML.

### Metformin inhibits WWP1 to enhance RUNX1-mediated apoptosis and impede CML progression both in vitro and in vivo

Since metformin exhibited an anti-CML mechanism from the above experiments [32]. To investigate the therapeutic significance of CML, we treated K562 and KCL22 cells with metformin or compound C. Immunoblot analysis revealed that metformin treatment reduced WWP1 expression and increased RUNX1 levels. The molecular change was correlated with heightened expression of pro-apoptotic markers and a concomitant decrease in anti-apoptotic proteins, indicating activation of apoptotic signaling (Figure 7A and Supplementary Figures 8A and 8 B). To corroborate these findings, apoptotic cell death was measured using the DAD(R) apoptosis detection kit. Cells subjected to metformin treatment demonstrated 50.21% apoptotic cell death compared to the control group (4.77%). The pro-apoptotic effect was counteracted by compound C treatment, suggesting that AMPK activation is the principal mechanism involved (Figure 7B). Consistent with these findings, a soft agar colony formation assay demonstrated significantly reduced colony growth in the metformin-treated cells, highlighting its inhibitory effect on leukemic cell proliferation (Figure 7C-F).

**Figure 7:**
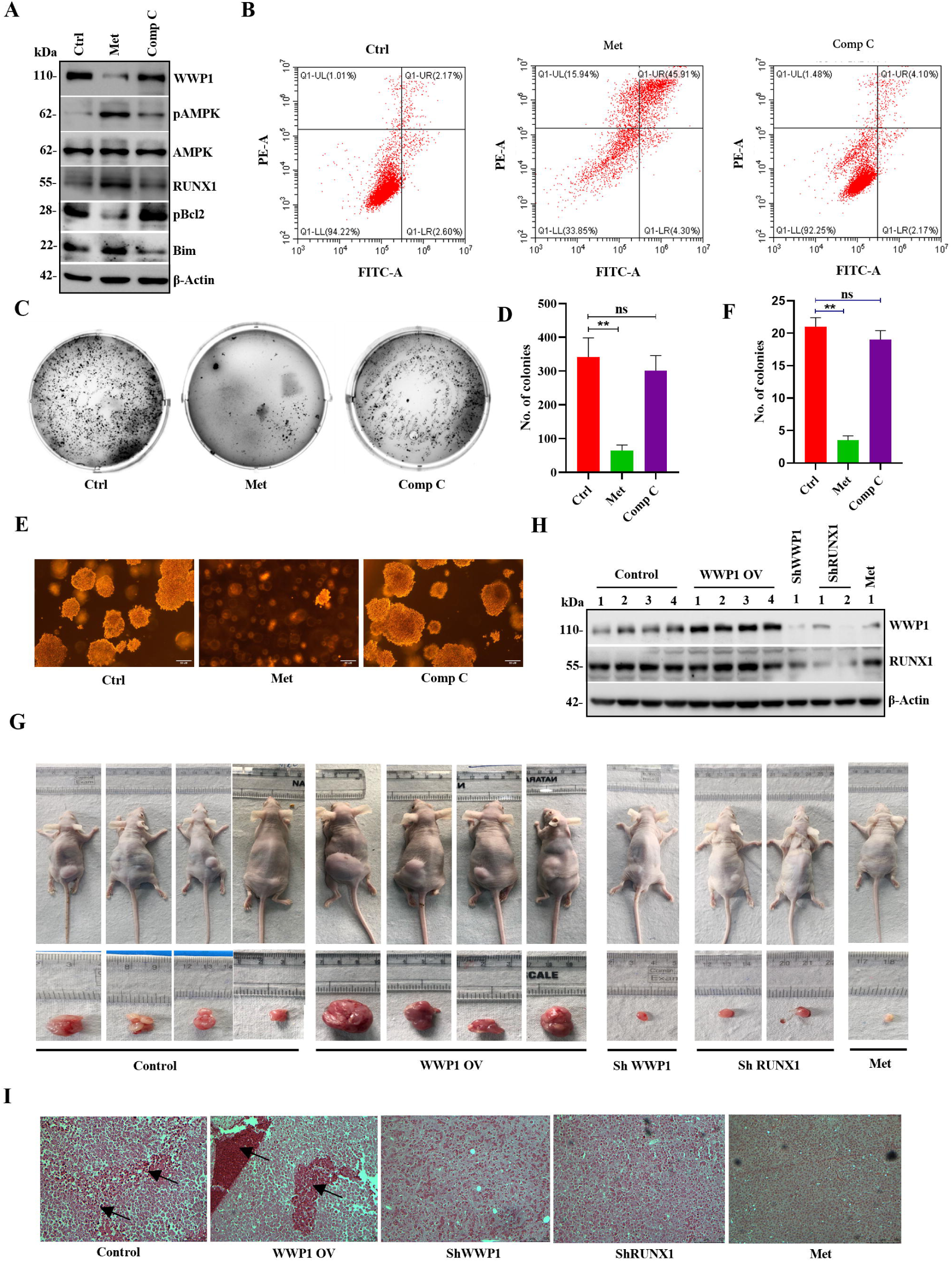
Metformin suppresses WWP1 signaling to induce apoptosis and inhibit CML progression *in vitro* and *in vivo*. **A.** Immunoblot analysis of K562 cells treated with metformin (20 mM, 24 h) and compound C (5 μM, 6 h). Whole-cell lysates were analyzed using the indicated antibodies. **B.** Flow cytometric analysis of apoptosis in K562 cells treated with metformin (20 mM, 24 h) and compound C (5 μM, 5 h), followed by DAD/PI staining showing the percentage of apoptosis. **C.** Representative images of colony formation assay showing the clonogenic potential of K562 cells following treatment with metformin (20 mM, 24 h) and compound C (5 μM, 5 h). **D.** Quantification of colony numbers from the colony formation assay shown in (C). **E.** Representative bright-field images of colonies captured at higher magnification showing colony morphology and size following the treatment with metformin (20 mM, 24 h) and compound C (5 μM, 5 h). Scale bar, 50 μm. **F.** Quantification of colony numbers from the representative image shown in (E). **G.** Representative images of FOXN1 athymic nude mice bearing subcutaneous K562 xenografts expressing shCtrl, shWWP1, shRUNX1, or shWWP1+WWP1, together with the metformin-treated group. Excised tumors were collected at the experimental endpoint. **H.** Immunoblot analysis of xenograft tumor lysates using the indicated antibodies to examine the expression of WWP1 and RUNX1 proteins. **I.** Representative hematoxylin and eosin (H&E)-stained sections of xenograft tumors from the indicated treatment groups. The sections were carefully obtained from the central region of the tumor mass to ensure that the analysis represents tumor-specific histopathological changes, rather than those of the adjacent normal stroma. Scale bar: 100 μm. Data are presented as mean ± SEM from three independent biological experiments (n=3). In panels D and F, colony numbers in the metformin and compound C-treated groups were compared with the corresponding untreated control using one-way ANOVA followed by Tukey’s multiple comparison test. Statistical significance is indicated as **p<0.01; ns: not significant.

To validate these findings *in vivo*, a xenograft tumor model was established by subcutaneously implanting K562 cells into nude mice. Mice were classified into five categories: control, WWP1 overexpression, WWP1 knockdown, RUNX1 knockdown, and metformin-treated. Metformin was administered daily via oral gavage at a dosage of 300 mg/kg to the treatment group. After four weeks, tumors in the WWP1 overexpression group exhibited a significantly greater volume than in the controls, whereas both the WWP1 and RUNX1 knockdown groups showed reduced tumor size. Notably, metformin administration recapitulated the tumor-suppressive effects observed in the knockdown groups, resulting in a substantial reduction in tumor volume (Figure 7G).

The tumors were excised after 4 weeks and subjected to immunoblot analysis and immunohistochemistry with hematoxylin and eosin (H&E) staining. Immunoblot analysis showed elevated levels of both WWP1 and RUNX1 in larger tumors, whereas metformin-treated tumors showed reduced WWP1 expression and increased RUNX1 levels, thus correlating with our in vitro results (Figure 7H). Histopathological evaluation revealed substantial leukemic cell infiltration and sinusoidal collapse in control tumors. Tissues with elevated WWP1 expression had extensive infiltration of undifferentiated cells. In contrast, tumors from the WWP1- or RUNX1-knockdown cohorts exhibited reduced leukemic load and improved tissue architecture. Tumors isolated from metformin-treated mice exhibited near-normal histology, with minimal leukemic infiltration and preserved cellular organization (Figure 7I). These findings collectively show that metformin promotes AMPK activation, leading to WWP1 suppression and RUNX1 stability.

## Discussion

CML remains a major clinical challenge, particularly in adult patients. Understanding the molecular mechanisms that drive its progression is crucial for developing effective strategies. RUNX1, a key transcription factor in hematopoiesis, has been implicated in several hematological malignancies, yet its regulation in CML remains poorly understood. In this study, we have demonstrated a novel regulatory mechanism for RUNX1 stability through post-translational modifications, such as ubiquitination facilitated by the E3 ubiquitin ligase WWP1 and phosphorylation by one of the metabolic regulators, AMPK.

Our findings revealed a significant correlation between WWP1 and RUNX1 protein levels in leukemic cell lines, further supported by in vivo experimental and clinical data. We show that E3 ubiquitin ligase WWP1 polyubiquitinates RUNX1 via K63-linkage. This ubiquitination helps stabilize RUNX1 and promote its nuclear localization, thereby facilitating the transcription of its downstream survival genes. These findings align with our earlier observations that RUNX1 is phosphorylated and maintained by AMPK [23], establishing a dual regulatory axis in which RUNX1 undergoes both ubiquitination and phosphorylation, exerting distinct mechanisms in CML progression (Figure 8).

**Figure 8:**
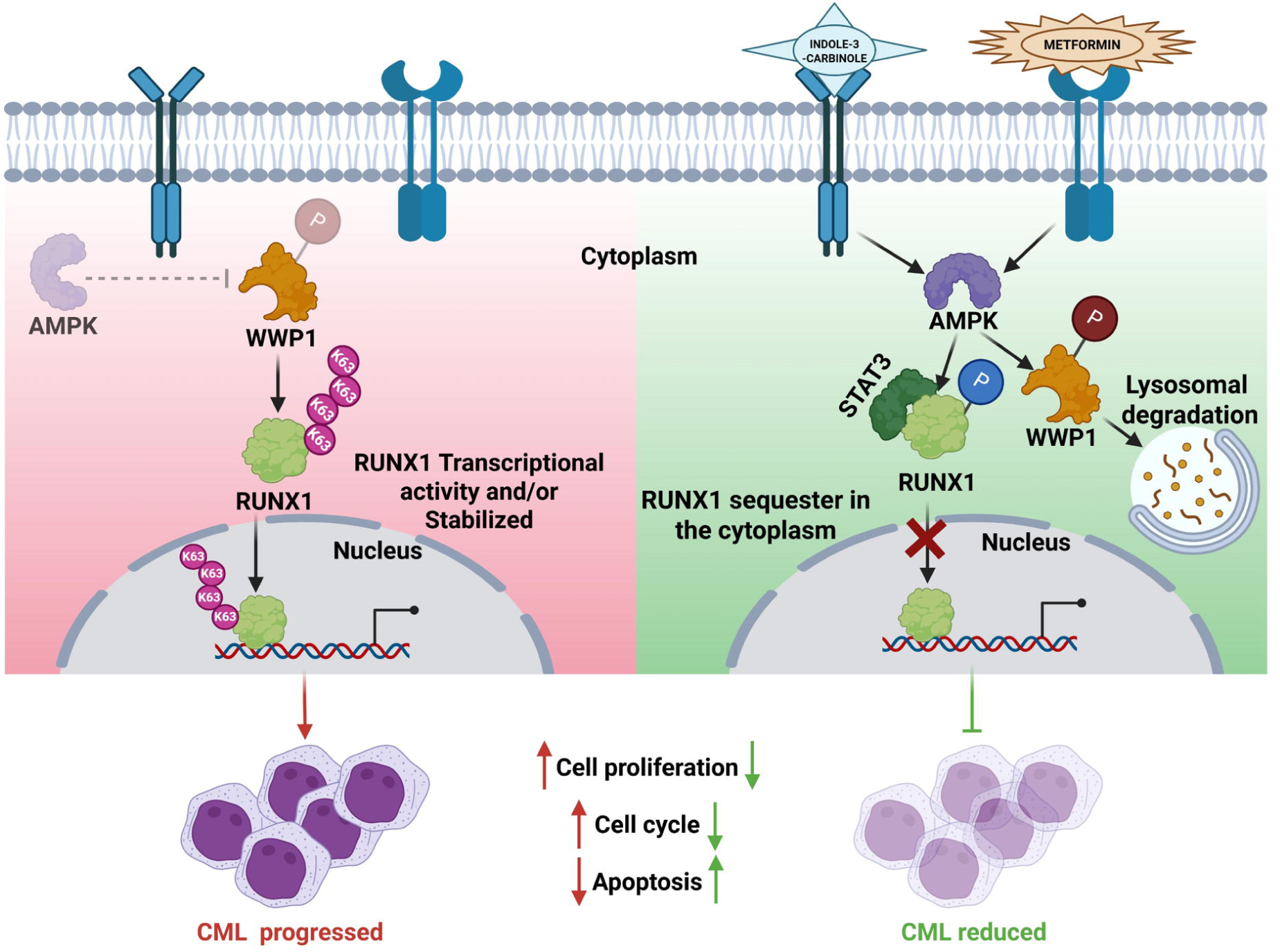
Schematic model illustrating WWP1 and AMPK-mediated differential regulation of RUNX1 in CML progression. The schematic representation depicts two distinct regulatory pathways: **Left panel** - Under basal conditions, WWP1 catalyses K63-linked polyubiquitination of RUNX1, promoting its stabilization and nuclear localization, where RUNX1 activates the transcription of genes involved in leukemic cell proliferation, cell-cycle progression, and survival, thereby contributing to CML progression. **Right panel** – Upon metformin-mediated activation of AMPK, WWP1 is phosphorylated at Thr692, triggering its lysosomal degradation and reducing WWP1-mediated K63-linked ubiquitination of RUNX1. Concurrently, AMPK phosphorylates RUNX1 at Ser94, promoting its association with STAT3 and cytoplasmic sequestration, thereby preventing RUNX1 nuclear translocation and transcriptional activity. Collectively, activation of the AAMPK-WWP1-RUNX1 signaling axis supresses leukemic cell proliferation, induces apoptosis, and inhibits CML progression.

Prior studies have recognized E3 ubiquitin ligases, including APC [10], RNF38 [11], CHIP [12], and STUB1 [13] as regulators of RUNX1 during development and erythroid differentiation. WWP1, a member of the HECT-type E3 ubiquitin ligase family, is recognized for its diverse functions in both physiological and pathological situations, including carcinogenesis, where it targets numerous substrates such as oncogenes and tumor suppressors. Notably, its overexpression is associated with a poor prognosis in several solid malignancies [15]. Our findings extend WWP1’s significance in leukemia by demonstrating its interaction with RUNX1. This interaction’s functional significance is shown by WWP1’s control of RUNX2 in bone biology [33]. Mechanistically, WWP1-facilitated K63-linked polyubiquitination of RUNX1 augments its intracellular localization and stimulates CML cell proliferation. Recent evidence in AML indicates that JARID1B is a substrate of WWP1, which improves chemoresistance and DNA damage repair capability by facilitating K63-mediated polyubiquitination of JARID1B [16]. Together, these findings suggest a larger paradigm in which WWP1-mediated K63 ubiquitination acts as a non-proteolytic oncogenic regulatory mechanism in myeloid malignancies. A similar mechanism has been observed with ΔNp63 in keratinocytes [34] and p53 stability modulation [35]. In addition, WWP1 has been implicated in AML by downregulating p27^Kip1^, thereby driving cell cycle progression and inhibiting autophagy signaling [17]. In WWP1-depleted AML cells, TXNIP-induced oxidative stress and DNA damage contribute to leukemic cell death [18]. Consistent with these reports, we observed that depletion of WWP1 in CML cells induced G2/M arrest, accumulation of DNA damage, and apoptosis. RUNX1 levels typically peak at G2/M transition and are degraded at the G2/M–G1 boundary, processes often involving SCF or APC ligases [7, 10]. Thus, WWP1 appears to reinforce cell-cycle–associated stabilization of RUNX1, promoting leukemic growth.

Ubiquitination is a central determinant of protein turnover and function, and cross-talk with phosphorylation further diversifies its regulatory outcomes [36, 37]. We previously demonstrated that AMPK phosphorylates RUNX1 at S94, promoting cytoplasmic sequestration via STAT3 interaction and thereby reducing leukemic potential [20, 22]. Although HECT-type E3 ubiquitin ligases are constitutively active in cancer cells, emerging evidence indicates that they are tightly regulated by phosphorylation [19, 38, 39]. In the present study, we identify WWP1 as a novel substrate of AMPK. Specifically, AMPK phosphorylates WWP1 at Thr692, triggering its lysosomal degradation. Pharmacological activation of AMPK by metformin reduced WWP1 levels and concurrently stabilized RUNX1 in CML primary cell lines. Similar effects were observed with indole-3-carbinol (I3C), a natural phytochemical with AMPK-activating and antitumor properties [26, 31]. Interestingly, AMPK-driven WWP1 degradation may involve recruitment of CYRR1, a cysteine- and tyrosine-rich membrane protein implicated in K63-linked lysosomal degradation and breast cancer survival [40]. This suggests that elevated glucose levels in skeletal muscle cells [41] could modulate WWP1 stability through AMPK-CYRR1-lysosomal pathways. These findings align with studies suggesting that K63-linked ubiquitination serves as a critical mechanism in transcriptional and metabolic regulation [42]. Similar mechanisms were also shown with SKP2-facilitated AKT ubiquitination, which promotes HK2 mitochondrial localization and cancer progression [36].

Our research underscores the therapeutic promise of targeting metformin, a widely used antidiabetic molecule, alone or using a combination of WWP1 inhibitor, I3C, a dietary phytochemical, by modulating the WWP1-AMPK-RUNX1 axis as a prospective therapeutic approach in CML [32, 43]. Our findings provide a foundation for the preclinical assessment of metformin and I3C in CML, to be subsequently validated in larger patient cohorts and clinical trials [44–46]. In addition to repurposing existing drugs, our findings suggest that WWP1 could be a viable target for PROTAC-mediated degradation, offering an opportunity to refine RUNX1 control and enhance therapeutic outcomes.

## Declarations

### Ethical approvals & Consent to participate

The study was approved by the NIMS Institutional Ethics Committee of Nizam’s Institute of Medical Sciences, Panjagutta, Hyderabad, Telangana, India, 500082, approval number (EC/NIMS/2350/2019). Human blood samples were collected only after obtaining written informed consent from all participants. The study was conducted in accordance with the approved institutional protocol and the ethical principles of the Department of Helsinki, in accordance with national guidelines for biomedical research involving human participants. Participants’ confidentiality and privacy were strictly maintained throughout the study, and all data were anonymized prior to analysis and publication.

## Consent for publication

All the authors read the final draft and approved it for publication.

## Data Availability

All the data is available in the main text or in the supplementary material as stated in the text.

## Authorship Contributions

AB Conceptualised, executed and analysed experiments, prepared and revised manuscript; SCG, MER, NV analysed experiments; DP, SG provided critical research material; RKK conceptualised, research material, manuscript editing; ABMR Conceptualised, analysed the data, secured funding, monitored the project and revised the manuscript.

## Acknowledgments

The University of Hyderabad funded the laboratory through institutional funding from DST-FIST-Level II, Institute of Eminence (IoE) (UoH-IoE-IIRC-23-001), MoE-SPARC/2019-2020/P2288/SL, DBT-SAHAJ/BUILDER (BT/INF/22/SP41176/2020), and ANRF-PAIR(ANRF/PAIR/2025/000012/EPAIR). ABMR acknowledges support from DST-SERB (EMR/2016/007595) and DBT-Genome Engineering Technologies (BT/PR38397/GET/119/306/2020), Government of India, as independent extramural grants. AB acknowledges the ICMR-SRF fellowship (ICMR-SRF No.2021-13931/CMB-BMS) for doctoral studies. AB acknowledges Dr. Surya Singh, Ms. Sanjibani Bag, and Mr. Kartikeya Saxena for the experimental insights. The authors acknowledge that ChatGPT (OpenAI) was used to refine the language and grammar in the editing of the text on some occasions. Images were developed using BioRender.com software. All scientific concepts, study design, data analysis, interpretation of results, and conclusions were independently developed, verified, and approved by the authors.

## Figure Legends

**Supplementary figure 1: WWP1 and RUNX1 expression in leukemic cell lines. A.** Densitometric quantification of WWP1 and RUNX1 expression in the indicated leukemic cell lines, with B cells used as the control. Data are presented as mean ± SEM from three independent biological experiments (n=3). In panel A, the relative protein expression of WWP1 and RUNX1 in each leukemia cell line was individually compared with B-cells. Statistical significance was determined using an unpaired, two-tailed Student’s t-test. Statistical significance is indicated as *p<0.05, **p<0.01. **B.** Expression analysis using GEPIA showing significantly elevated WWP1 and RUNX1 expression in AML patient samples (*n* = 173) compared with healthy individuals (*n* = 70). Expression values are presented as transcripts per million (TPM) and were quantified using RSEM with log2 transformation (pseudo-count = 1) for normalization. **C.** Pearson correlation analysis demonstrating a strong positive correlation between WWP1 and RUNX1 expression in AML samples (*R* = 0.82). **D&E.** Kaplan– Meier survival analyses showing the association between WWP1 (D) and RUNX1 (E) expression and overall survival in AML patients. Patients were stratified according to high and low expression levels of the respective genes.

**Supplementary figure 2: WWP1 interacts with and regulates RUNX1 protein stability:** WWP1 interacts with and regulates RUNX1 protein stability in leukemic cell lines. **A.** Protein sequence analysis of RUNX1 identified a PPPY-containing motif, suggesting a potential docking site for the WW domain of WWP1. **B&C.** Reciprocal co-immunoprecipitation (co-IP) assays in KCL22 cells using anti-WWP1 and anti-RUNX1 antibodies demonstrate an endogenous physical interaction between WWP1 and RUNX1. **D&E.** Reciprocal co-IP assays in Jurkat J6 cells confirm the endogenous interaction between WWP1 and RUNX1. **F&G.** Reciprocal co-IP assays in HL-60 cells further confirm the endogenous interaction between WWP1 and RUNX1. **H-J.** Protein–protein docking analysis using the HDOCK server predicts an interaction between the WW3 domain of WWP1 and the PPPY motif of RUNX1, with the docking model indicating a potential salt-bridge interaction between WW3 domain of WWP1 and PPPY motif of RUNX1. **K&L.** WWP1 was knocked down in K562 and KCL22 cells using shRNA, and WWP1 protein levels were assessed by western blotting. **M.** Densitometric quantification demonstrates a significant reduction in WWP1 protein levels following WWP1 knockdown (P < 0.01), accompanied by a significant reduction in RUNX1 protein levels (P < 0.05). **N.** Densitometric quantification of WWP1 and RUNX1 protein levels following dose-dependent ectopic expression of Myc-tagged WWP1 in K562 cells demonstrates a progressive increase in WWP1 and RUNX1 protein levels, with statistically significant differences indicated by (P < 0.05) and (P < 0.01). **O.** Quantitative real-time PCR analysis demonstrates a significant increase in WWP1 mRNA following WWP1 overexpression (P < 0.001), whereas RUNX1 mRNA levels are not significantly altered. Data are presented as mean ± SEM. Statistical significance was determined using an unpaired, two-tailed Student’s t-test. Statistical significance is indicated as *p < 0.05, **p < 0.01, and ***p < 0.001; ns, not significant.

**Supplementary figure 3: WWP1 promotes RUNX1 protein stability through K63-linked polyubiquitination. A.** *In silico* prediction of potential RUNX1 E3 ubiquitin ligases using UbiBrowser, with the predicted confidence scores shown. WWP1 was identified as a high-confidence candidate regulator of RUNX1 ubiquitination. **B.** K562 cells treated with I3C were analysed for endogenous protein ubiquitination by immunoblotting with an anti-ubiquitin (Ub) antibody; WWP1, RUNX1 and β-actin were assessed in parallel. **C-E.** Endogenous protein ubiquitination and WWP1 and RUNX1 protein expression were analysed in KCL22 (**C**), Jurkat J6 (**D**), and HL-60 (**E**) cells following WWP1 depletion with shRNA. F & **G.** Jurkat J6 (**F**) and HL-60 (**G**) cells treated with I3C were analysed for endogenous protein ubiquitination and WWP1 and RUNX1 protein expression by immunoblotting. **H.** HEK293T cells were transfected with the indicated GFP-RUNX1, Myc-WWP1, or catalytically inactive Myc-WWP1 C890A constructs together with HA-ubiquitin. Cells were lysed under denaturing conditions, and ubiquitinated proteins were immunoprecipitated using an anti-HA antibody and analysed by immunoblotting for RUNX1. WWP1 and RUNX1 expression in the input lysates was assessed as indicated. **I.** Quantification of RUNX1 protein turnover in HEK293T cells transiently expressing vector, WWP1, or WWP1 C890A. Cells were treated with cycloheximide (CHX; 100 μg/mL) for the indicated times, and RUNX1 protein abundance was determined by immunoblotting and expressed relative to the 0 h level. Data are presented as mean ± SEM from three independent biological experiments (n=3). RUNX1 protein levels at each CHX treatment time point in WWP1 and WWP1C890A cells were compared with the corresponding vector control using two-way ANOVA followed by Sidak’s multiple comparisons test. Statistical significance is indicated as *p<0.05, **p<0.01.

**Supplementary figure 4: WWP1 regulates CML cell growth and survival through RUNX1. A.** Representative bright-field images (20X) of colony formation assays in K562 cells expressing control shRNA or WWP1-targeting shRNA. **B.** Quantification of colony numbers from six randomly selected fields per condition. **C.** Immunoblot analysis of KCL22 cells expressing shCtrl or shWWP1 showing the indicated proteins involved in DNA damage response and cell-cycle regulation. **D.** Flow cytometric analysis of DAD/PI-stained K562 cells expressing shCtrl or shWWP1, showing increased apoptosis following WWP1 depletion. **E.** Immunoblot analysis of whole-cell lysates from KCL22 cells expressing shCtrl or shWWP1 for the indicated proteins associated with cell-cycle progression and apoptosis. **F&G.** Cell proliferation of K562 (**F**) and KCL22 (**G**) cells stably expressing shCtrl, shWWP1, shRUNX1, shWWP1 together with WWP1 re-expression, or shWWP1 together with RUNX1 re-expression, as assessed by PrestoBlue assay over the indicated time points. **H.** Quantification of colony numbers from the colony formation assay shown in Fig. 3K in K562 cells expressing the indicated shRNAs and rescue constructs. **I.** Representative images of colony formation assays in KCL22 cells expressing shCtrl, shWWP1, shRUNX1, shWWP1 together with WWP1 re-expression, or shWWP1 together with RUNX1 re-expression. **J.** Quantification of colony numbers from the colony formation assay shown in (I). **K.** Immunoblot analysis of KCL22 cells following WWP1 and/or RUNX1 depletion and re-expression, showing the indicated proteins involved in cell-cycle progression and apoptosis. **L.** Densitometric quantification of WWP1, RUNX1, Cyclin B1, pBcl2 and Bim protein expression in KCL22 cells under the indicated experimental conditions. β-Actin was used as the loading control. Data are presented as mean ± SEM. For comparisons between two groups (**B**), statistical significance was determined using a two-tailed unpaired Student’s *t*-test. For comparisons among multiple groups (**H, J and L**), statistical significance was determined by one-way ANOVA followed by Tukey’s multiple-comparisons test. For **L**, statistical analysis was performed separately for each indicated protein. Statistical significance is indicated as *p<0.05, **p<0.01, ***p<0.001; ns: not significant.

**Supplementary figure 5: WWP1 depletion delays cell-cycle progression through G2/M in K562 cells. A&B.** Cell-cycle distribution of synchronized K562 cells expressing control shRNA (shCtrl; A) or WWP1-targeting shRNA (shWWP1; B). Cells were synchronized by sequential thymidine and nocodazole treatment and subsequently released for the indicated time periods. DNA content was analysed by flow cytometry, and the percentages of cells in G1, S and G2/M phases are indicated. WWP1 depletion resulted in sustained accumulation of cells in the G2/M phase following release from nocodazole synchronization, consistent with delayed G2/M progression or exit.

**Supplementary figure 6: AMPK negatively regulates WWP1 protein abundance through phosphorylation. A.** Schematic representation of WWP1 and the WWP1 phosphorylation-site mutants generated to investigate AMPK-dependent phosphorylation. The indicated Alanine (A) and aspartate (D) substitutions were introduced at the predicted AMPK phosphorylation site. **B.** Dose- and time-dependent effects of metformin on K562 cell viability, assessed at the indicated concentrations and time points. **C.** Densitometric quantification of WWP1, pAMPK, total AMPK, and RUNX1 protein expression in K562 cells treated with metformin at the indicated time points. **D.** Immunoblot analysis of WWP1, pAMPK, total AMPK and RUNX1 expression in KCL22 cells treated with increasing concentrations of metformin for 24 h. **E.** Densitometric quantification of the indicated proteins in KCL22 cells treated with metformin. **F.** RT-PCR analysis of WWP1 and RUNX1 mRNA expression in K562 cells treated with metformin (20 mM), showing that metformin treatment does not significantly alter WWP1 transcript abundance. **G.** Dose- and time-dependent effects of I3C on K562 cell viability. **H.** Immunoblot analysis of WWP1, pAMPK and RUNX1 protein expression in K562 cells treated with increasing concentrations of I3C for 24 h. **I.** Densitometric quantification of WWP1, pAMPK and RUNX1 protein expression following I3C treatment. **J.** Immunoprecipitation analysis of K562 cell lysates treated with metformin (20 mM). WWP1 was immunoprecipitated and analysed by immunoblotting with antibodies against pAMPK and AMPK substrate motifs. For (B, G), data are presented as mean ± SEM, and statistical significance was determined by two-way ANOVA, with concentration and treatment duration as the respective factors, followed by Dunnett’s multiple-comparisons test where indicated. For (C, E, I), statistical significance was determined using one-way ANOVA followed by Dunnett’s multiple comparisons test, with each treatment condition compared with its corresponding untreated control. For (F), statistical significance was determined using a two-tailed unpaired Student’s *t*-test. Statistical significance is indicated as *p<0.05, **p<0.01; ns: not significant.

**Supplementary figure 7: AMPK activation regulates WWP1 and RUNX1 protein abundance. A.** HEK293T cells transiently expressing the indicated WWP1 and/or RUNX1 constructs were treated with metformin (20 mM) for 24 h, and whole-cell lysates were analysed by immunoblotting for WWP1, RUNX1, pAMPK and total AMPK. β-Actin was used as a loading control. **B.** Densitometric quantification of WWP1, pAMPK, total AMPK and RUNX1 protein levels from the immunoblots shown in (A). Protein abundance was normalized to β-actin and expressed relative to the corresponding control condition. **C.** HEK293T cells transfected with the indicated constructs were treated with metformin (20 mM, 24 h) and/or the proteasome inhibitor MG132 (1mg/ml, 6 h), as indicated. Whole-cell lysates were analysed by immunoblotting for WWP1, pAMPK and total AMPK to assess the contribution of proteasomal degradation to metformin-mediated regulation of WWP1. Data are presented as mean ± SEM. For (B), statistical significance was determined separately for each protein by one-way ANOVA followed by Dunnett’s multiple-comparisons test, with the indicated treatment groups compared with the corresponding control condition. Statistical significance is indicated as *p<0.05, **p<0.01, ***p<0.001; ns: not significant.

**Supplementary figure 8: AMPK-dependent inhibition of WWP1 is associated with apoptotic signaling in KCL22 cells. A.** KCL22 cells were treated with metformin (Met; 20 mM, 24 h) or the AMPK inhibitor Compound C (Comp C; 5 μM, 6 h), and whole-cell lysates were analysed by immunoblotting for WWP1, RUNX1, pAMPK, total AMPK, pBCL2 and Bim. β-Actin was used as a loading control. **B.** Densitometric quantification of the indicated proteins from the immunoblots shown in (A). Protein expression was normalized to β-actin and expressed relative to the untreated control. Data are presented as mean ± SEM. Statistical significance was determined separately for each protein by one-way ANOVA followed by Dunnett’s multiple-comparisons test, with metformin- and Compound C-treated groups compared with the untreated control. Statistical significance is indicated as *p<0.05, **p<0.01; ns: not significant.

